# Relating scene memory and perception activity to functional properties, networks, and landmarks of posterior cerebral cortex - a probabilistic atlas

**DOI:** 10.1101/2025.01.06.631538

**Authors:** Adam Steel, Deepa Prasad, Brenda D. Garcia, Caroline E. Robertson

## Abstract

Adaptive behavior in complex environments requires integrating visual perception with memory of our spatial environment. Recent work has implicated three brain areas in posterior cerebral cortex — the place memory areas (PMAs) that are anterior to the three visual scene perception areas (SPAs) – in this function. However, PMAs’ relationship to the broader cortical hierarchy remains unclear due to limited group-level characterization. Here, we examined the PMA and SPA locations across three fMRI datasets (44 participants, 29 female). SPAs were identified using a standard visual localizer where participants viewed scenes versus faces. PMAs were identified by contrasting activity when participants recalled personally familiar places versus familiar faces (Datasets 1-2) or places versus multiple categories (familiar faces, bodies, and objects, and famous faces; Dataset 3). Across datasets, the PMAs were located anterior to the SPAs on the ventral and lateral cortical surfaces. The anterior displacement between PMAs and SPAs was highly reproducible. Compared to public atlases, the PMAs fell at the boundary between externally-oriented networks (dorsal attention) and internally-oriented networks (default mode). Additionally, while SPAs overlapped with retinotopic maps, the PMAs were consistently located anterior to mapped visual cortex. These results establish the anatomical position of the PMAs at inflection points along the cortical hierarchy between unimodal sensory and transmodal, apical regions, which informs broader theories of how the brain integrates perception and memory for scenes. We have released probabilistic parcels of these regions to facilitate future research into their roles in spatial cognition.

**Significance statement:** Complex behavior requires the dynamic interplay between mnemonic and perceptual information. For example, navigation requires representation of the current visual scene and its relationship to the surrounding visuospatial context. We have suggested that the place memory areas, three brain areas located anterior to the scene perception areas in visual cortex, are well-positioned to serve this role. Here, in a large group of participants, we show that the place memory areas are robustly localizable, and that their position at the interface of multiple distributed brain networks is uniquely suited to mnemonic-perceptual integration. We have released probabilistic regions-of-interest and localization procedure so that others can identify these areas in their own participants.

## Introduction

The integration of perception and memory is a fundamental challenge for the brain, requiring neural systems to simultaneously maintain current sensory input while accessing relevant stored information(Rust and Palmer, 2021). This integration is particularly evident during spatial navigation, where we dynamically exchange information between the immediate visual scene and memory of the broader environment to interpret visible features, predict unseen elements, and guide attentional and motor decisions(Robertson et al., 2016; Brunec et al., 2018; Haskins et al., 2020; Berens et al., 2021; Draschkow et al., 2022). Understanding how the brain’s functional architecture enables perceptual and mnemonic representations to effectively interface, while also avoiding interference, is therefore a central question in cognitive neuroscience(Summerfield and De Lange, 2014; Kiyonaga et al., 2017; Favila et al., 2020; Libby and Buschman, 2021).

Recent work has identified a network of three brain areas in posterior cerebral cortex that may play a crucial role in bridging perception and memory in the domain of scenes(Steel et al., 2021, 2023, 2024b). These “place memory areas”, on the lateral, ventral, and medial surfaces (LPMA, VPMA, MPMA, respectively), selectively activate when individuals recall familiar places (e.g., their house) compared to other memorable stimuli like familiar people (e.g., their mother) (Steel et al., 2021). Critically, each place memory area is located immediately anterior and adjacent to one of the three functionally-defined “scene perception areas” in high-level visual cortex - the occipital place area (OPA), parahippocampal place area (PPA), and medial place area (MPA, also known as retrosplenial complex), on the brain’s lateral, ventral, and medial surfaces, respectively (Epstein and Kanwisher, 1998; Hasson et al., 2003; Epstein et al., 2007; Dilks et al., 2013; Silson et al., 2016). This systematic topographical relationship suggests the place memory areas are anatomically positioned to directly access perceptual representations of scenes(Steel et al., 2023), which may change with age (Srokova et al., 2022).

Functionally, the place memory areas exhibit several key properties that highlight their role in bridging perception and memory. The PMAs areas contain multivariate representations of specific remembered scene views(Bainbridge et al., 2020; Steel et al., 2023), and activation in these areas scales with the amount of remembered visuospatial context associated with a viewed scene (Steel et al., 2023). Connectivity analyses show that the PMAs constitute a distinct functional network from the SPAs, which is more strongly connected with spatial memory structures like the hippocampus compared to early visual cortex (Baldassano et al., 2016; Silson et al., 2016, 2019; Steel et al., 2021). Additionally, PMAs are dynamically connected to SPAs through voxel-wise retinotopically opponent patterns during tasks requiring perceptual-mnemonic interaction, such as recognizing familiar scenes (Steel et al., 2024b, 2024a). Finally, the anterior shift of memory compared with perception may be specific to scene stimuli as compared with faces (Steel et al., 2021; Chen et al., 2024). Taken together, this organization aligns with a specialized role in processing out-of-view spatial information during navigation (Steel et al., 2023).

While these properties make the place memory areas compelling candidates for integrating perceptual and mnemonic representations of scenes, their broader relationship to large-scale cortical topography remains unclear. This gap stems partly from methodological choices in prior research. Given the individualized functional location of category-selective areas in high-level visual cortex (Kanwisher et al., 1997; Epstein and Kanwisher, 1998; Grill-Spector and Weiner, 2014; Kanwisher, 2017), most investigations have employed subject-specific localization approaches to define these areas (Steel et al., 2021, 2023, 2024b; Srokova et al., 2022), leaving open questions about whether the PMAs represent a fundamental feature of human cortical organization or whether their location varies substantially across individuals and methodological approaches. Additionally, because group-level analyses have been limited, the place memory areas’ relationships with other known anatomical and functional landmarks (e.g., retinotopic maps(Wandell et al., 2007; Wang et al., 2015) or large-scale cortical networks (Raichle et al., 2001; Fox et al., 2005; Thomas Yeo et al., 2011; Yeo et al., 2015; Braga and Buckner, 2017; DiNicola et al., 2020; Du et al., 2024) and functional gradients(Margulies et al., 2016; Huntenburg et al., 2018; Reznik et al., 2024)) remain poorly characterized. Understanding these relationships can provide important insight into the role of these memory areas in the brain outside of the visual system and shed light on the relationship between spatial memory processes and broader large-scale organizing principles of the human brain.

Here we address these open questions by characterizing the topography of place memory areas across three independent datasets that vary in their fMRI acquisition parameters, preprocessing approaches, and experimental designs. This approach allows us to establish the consistency of these areas’ locations relative to scene perception areas and other cortical landmarks, while controlling for methodology-specific effects. Our results suggest the place memory areas – each located immediately anterior to a classic scene perception area on the ventral and dorsal surface of the human brain – represent a robust and consistent feature of human functional cortical organization. Further, we establish their anatomical position at inflection points along the cortical hierarchy between unimodal sensory and transmodal, apical regions, which informs broader theories of how the brain integrates perception and memory. Finally, we have made probabilistic parcels defining their locations freely available to the research community to facilitate future research on these areas, and to better understand their probabilistic relationship with reference to other probabilistic atlases of functional areas in the human visual system (Julian et al., 2012; Wang et al., 2015; Rosenke et al., 2021).

## Methods

### Procedure

The data for this study comprised functional localizers from three independent datasets. Dataset 1 and Dataset 3 shared 3 participants, otherwise, all participants were non-overlapping. Dataset 1 (Steel et al., 2021)and Dataset 2 (Steel et al., 2022, 2023, 2024b)were analyzed for different studies that have been previously published. Dataset 3 was collected for a separate study that has not been published.

The datasets in this study differed on their imaging parameters and task paradigms, which enabled us to test the impact of these factors on the robustness of the place memory and scene perception area location. Datasets 1 and 2 had different imaging parameters but identical task paradigms. Datasets 2 and 3 had identical imaging parameters but different task paradigms. Datasets 1 and 3 had different imaging parameters and task paradigms. A general description of the relationships between these datasets is detailed in Table 1, and a detailed description of all relevant details are reported below.

**Table 1.** Relationship between Datasets 1-3.

| Dataset | Number of participants | MRI acquisition / processing | Task conditions (imagery) | Total data acquired (Perception task) | Total data acquired (Memory task) | Publications |
| --- | --- | --- | --- | --- | --- | --- |
| Dataset 1 | 14 | Single echo Standard | Familiar places<br>Familiar faces | 2 runs, 226 TRs (run duration 552 s) | 4 runs, 155 TRs (run duration 310 s) | Steel et al., 2021 |
| Dataset 2 | 23 | Multi-echo ME-ICA | Familiar places<br>Familiar faces | 2 runs, 226 TRs (run duration 552 s) | 4 runs, 155 TRs (run duration 310 s) | Steel et al., 2022;<br>Steel et al., 2023;<br>Steel*, Silson*, et al., 2024 |
| Dataset 3 | 10<br>(3 shared with Dataset 1) | Multi-echo ME-ICA | Familiar places<br>Familiar faces<br>Familiar bodies<br>Familiar objects<br>Famous faces | - | 6 runs, 234 TRs (run duration 768 s) | Unpublished |

### Participants

Data from 44 unique participants (Age: 24.9±6.1 s.d., 15 male) comprise this study and are distributed across three different datasets (Dataset 1: N = 14, Age: 25.6±4.1 s.d., 5 male; Dataset 2: N = 23, Age: 24.6±7.7 s.d., 8 male; Dataset 3: N =10, Age: 25.8±3.3 s.d., 4 male). Three participants were present in both Dataset 1 and Dataset 3. All subjects had normal or correct to normal vision, were not colorblind, and were free from neurological or psychiatric conditions. Written consent was obtained from all participants in accordance with the Declaration of Helsinki and with a protocol and consent form approved by the Dartmouth College Institutional Review Board (Protocol #31288). Participants were compensated for their time at a rate of $20/hr. No statistical methods were used to pre-determine sample sizes.

### Visual Stimuli and Tasks

#### Perception and memory tasks - Datasets 1 and 2

##### Static scene perception area localizer

The scene perception areas (SPAs, i.e. occipital place area, OPA; parahippocampal place area, PPA; medial place area, MPA) are regions that selectively activate when an individual perceives places (i.e., a kitchen) compared with other categories of visual stimuli (i.e., faces, objects, bodies) (Epstein and Kanwisher, 1998; Silson et al., 2016; Weiner et al., 2018; Steel et al., 2021). To identify these areas in each person, participants performed an independent functional localizer scan. On each run of the localizer (2 runs), participants passively viewed blocks of scene, face, and object images presented in rapid succession (500 ms stimulus, 500 ms ISI). Blocks were 24 s long, and each run comprised 12 blocks (4 blocks/condition). There was no interval between blocks.

##### Place memory area localizer

The place memory areas (PMAs) are defined as regions that selectively activate when a person recalls personally familiar places (i.e., their kitchen) compared with personally familiar people (i.e., their mother)(Steel et al., 2021). To identify these areas in each person, participants performed an independent functional localizer scan. Prior to fMRI scanning, participants generated a list of 36 personally familiar people and places to establish individualized stimuli (72 stimuli total). These stimuli were generated based on the following instructions.

> “For your scan, you will be asked to visualize people and places that are personally familiar to you. So, we need you to provide these lists for us. For personally familiar people, please choose people that you know in real life (no celebri>es) that you can visualize in great detail. You do not need to be contact with these people now, as long as you knew them personally and remember what they look like. So, you could choose a childhood friend even if you are no longer in touch with this person. Likewise, for personally familiar places, please list places that you have been to and can richly visualize. You should choose places that are personally relevant to you, so you should avoid choosing places that you have only been to one >me. You should not choose famous places where you have never been. You can choose places that span your whole life, so you could do your current kitchen, as well as the kitchen from your childhood home.”

During fMRI scanning, participants recalled these people and places. On each trial, participants saw the name of a person or place and recalled them in as much detail as possible for the duration that the name appeared on the screen (10 s). Trials were separated by a variable ISI (4-8 s). Place memory areas were localized by contrasting activity when participants recalled personally familiar places compared with people (see ROI definitions section). All trials were unique stimuli, and conditions (i.e., people or place stimuli) were pseudo-randomly intermixed so that no more than two repeats per condition occurred in a row.

#### Memory task - Dataset 3

Dataset 3 comprised two tasks, a multi-category dynamic perception task and a multi-category dynamic memory task. The data from the multi-category perception task is being reported in a separate publication and will not be discussed here.

##### Multi-category place memory localizer

Prior to fMRI scanning, participants generated a list of 5 examples from 5 categories: personally familiar people’s faces (e.g., wife’s face, sister’s face), familiar people’s body parts (e.g., mother’s hands, father’s feet), familiar places (e.g., my kitchen, college library), familiar objects roughly the size of a person’s face (e.g., laptop, rugby ball), and famous people’s faces (Beyonce’s face, Obama’s face) to establish individualized stimuli (72 stimuli total). These stimuli were generated based on the following instructions.

> “For your scan, you will be asked to visualize people’s faces and bodies, places, and objects that are familiar to you. So, we need you to provide these lists for us. For personally familiar people’s faces, please choose people that you know in real life (no celebri>es) that you can visualize in great detail. You do not need to be contact with these people now, as long as you knew them personally and remember what they look like. So, you could choose a childhood friend even if you are no longer in touch with this person.

> For bodies, you will choose 5 familiar parts of bodies that are specific to someone you know (e.g., the hands or feet of your parents). The parts of the body you choose do not have to belong to the personally familiar people you chose. Please do not include explicit body parts. Some examples of body parts specific to a person could be hands, feet, legs, arms, whole bodies etc., but not faces.

> For familiar objects, or objects, choose 5 personally familiar objects that are about the size of a toaster (e.g., your favorite mug, a co@ee maker). These objects should be personal items that you know very well and interact with fairly regularly (e.g., your personal hairbrush or your soccer ball). You should also be familiar with from multiple visual angles or viewpoints.

> For personally familiar places, please list places that you have been to and can richly visualize. You should choose places that are personally relevant to you, so you should avoid choosing places that you have only been to one >me. You should not choose famous places where you have never been. You can choose places that span your whole life, so you could do your current kitchen, as well as the kitchen from your childhood home.

> For famous people, list some celebri>es that you do not know personally whose faces you can visualize richly. You should be able to imagine their faces and bodies moving. For example, if you can imagine Beyonce’s face, then she would be a good choice. However, if you could only imagine Beyonce’s body while she is dancing, then you should pick a different celebrity.”

During fMRI scanning, participants performed 6 runs of the imagery task. During each run, participants recalled these people, body parts, objects, and places. All stimuli were presented in each run, and trials were pseudo-randomized so that no category could appear on more than 2 consecutive trials.

The trial sequence was modelled oa our prior work (Steel et al., 2023). On each trial, participants saw the name of a person or place and recalled them in as much detail as possible. The name of the stimulus was displayed for 1 second, followed by a 1 second dynamic mask (mix of ascii characters approximately the same length as the stimulus cue), followed by four circles (‘oooo’). Participants were instructed to maintain imagery for as long as the circles remained on the screen (10 s). Trials were separated by a variable ISI (4-8 s).

### MRI acquisition

#### T1-weighted anatomical scan

For registration purposes, a high-resolution T1-weighted magnetization-prepared rapid acquisition gradient echo (MPRAGE) imaging sequence was acquired (TR = 2300 ms, TE = 2.32 ms, inversion time = 933 ms, Flip angle = 8°, FOV = 256 × 256 mm, slices = 255, voxel size = 1 ×1 × 1 mm). T1 images were segmented and surfaces were generated using Freesurfer (Dale et al., 1999; Fischl et al., 2002; Fischl, 2012) (version 6.0) and aligned to the fMRI data using AFNI programs (Cox, 1996) align_epi_anat.py and @SUMA_AlignToExperiment(Saad and Reynolds, 2012).

#### Dataset 1 – Single Echo EPI

Single-echo T2*-weighted echo-planar images covering the temporal, parietal, and frontal cortices were acquired using the following parameters: TR=2000 ms, TE=32 ms, GRAPPA=2, Flip angle=75°, FOV=240 x 240 mm, Matrix size=80 x 80, slices=34, voxel size=3 x 3 x 3 mm. To minimize dropout caused by the ear canals, slices were oriented parallel to temporal lobe(Weiskopf et al., 2006). The initial two frames were discarded by the scanner to achieve steady state.

#### Dataset 2 & 3 – Multi Echo EPI

Multi-echo T2*-weighted sequence. The sequence parameters were: TR=2000 ms, TEs=[14.6, 32.84, 51.08], GRAPPA factor=2, Flip angle=70°, FOV=240 x 192 mm, Matrix size=90 x 72, slices=52, Multi-band factor=2, voxel size=2.7 mm isotropic. The initial two frames of data acquisition were discarded by the scanner to allow the signal to reach steady state.

### MRI Preprocessing

#### Dataset 1 – Single Echo EPI

Data was preprocessed using AFNI (version 20.3.02 ‘Vespasian’)(Cox, 1996). In addition to the frames discarded by the fMRI scanner during acquisition, the initial two frames were discarded to allow T1 signal to achieve steady state. Signal outliers were attenuated (3dDespike). Motion correction was applied, and parameters were stored for use as nuisance regressors (3dVolreg). Data were then iteratively smoothed to achieve a uniform smoothness of 5mm FWHM (3dBlurToFWHM).

#### Dataset 2 & 3 – Multi Echo EPI

Multi-echo data processing was implemented based on the multi-echo preprocessing pipeline from afni_proc.py in AFNI (version 21.3.10 ‘Trajan’)(Cox, 1996). Signal outliers in the data were attenuated (3dDespike(Jo et al., 2013)). Motion correction was calculated based on the second echo, and these alignment parameters were applied to all runs. The optimal combination of the three echoes was calculated, and the echoes were combined to form a single, optimally weighted time series (T2smap.py). Multi-echo ICA denoising(Kundu et al., 2012; Evans et al., 2015; DuPre et al., 2019, 2021) was then performed (see *Multi-echo ICA*, below). Following denoising, signals were normalized to percent signal change, and data were smoothed with a 3mm Gaussian kernel (3dBlurInMask).

##### Multi-echo ICA

The data were denoised using multi-echo ICA denoising (tedana.py(Kundu et al., 2012; Evans et al., 2015; DuPre et al., 2019; Steel et al., 2022), version 0.0.12). In brief, PCA was applied, and thermal noise was removed using the Kundu decision tree method. Subsequently, data were decomposed using ICA, and the resulting components were classified as signal and noise based on the known properties of the T2* signal decay of the BOLD signal versus noise. Components classified as noise were discarded, and the remaining components were recombined to construct the optimally combined, denoised timeseries.

### MRI analysis

#### Dataset 1 & 2 GLM and ROI definition

##### Scene perception area localizer

To define scene perceptual areas, the scene perception localizer was modeled by fitting gamma function of the block duration with a square wave for each condition (Scenes, Faces, and Objects) using 3dDeconvolve. Estimated motion parameters were included as additional regressors of no-interest along with 4^th^ order polynomials. Scene areas were drawn based on a general linear test comparing the coefficients of the GLM during scene versus face blocks. These contrast maps were then transferred to the SUMA standard mesh (std.141) using @SUMA_Make_Spec_FS and @Suma_AlignToExperiment.

Initial ROIs were hand drawn using a vertex-wise significance of p<0.001 (t-statistic > 3.3) along with expected anatomical locations (e.g., PPA was not permitted to extend posteriorly beyond the edge of the collateral sulcus, was used to define the pre-constrained regions of interest (Weiner et al., 2018; Steel et al., 2021).

##### Place memory area localizer

To define place memory areas, the familiar people/places memory data was modeled by fitting a gamma function of the trial duration for trials of each condition (people and places) using 3dDeconvolve. Estimated motion parameters were included as additional regressors of no-interest along with 4^th^ order polynomials. Activation maps were then transferred to the suma standard mesh (std.141) using @SUMA_Make_Spec_FS and @Suma_AlignToExperiment.

People and place memory areas were drawn based on a general linear test comparing coefficients of the GLM for people and place memory. Initial ROIs were hand drawn using a vertex-wise significance threshold of p<0.001 (t-statistic > 3.3). These ROIs were subsequently constrained to the top 800 most selective vertices before aggregating at the group level (see below)

#### Dataset 3 GLM and ROI definition

##### Dynamic place memory area localizer

To define place memory areas, the familiar people/places memory data was modeled by fitting a gamma function of the trial duration for trials of each condition (famous faces, familiar faces, familiar bodies, familiar objects, familiar places) using 3dDeconvolve (10 s). We included a single regressor to model the cue onset and stimulus mask for all trials (2 s). Estimated motion parameters were included as additional regressors of no-interest along with 4^th^ order polynomials. Activation maps were then transferred to the suma standard mesh (std.141) using @SUMA_Make_Spec_FS and @Suma_AlignToExperiment.

People and place memory areas were drawn based on a general linear test comparing coefficients of the GLM for place memory versus all other categories (personally familiar faces, body parts, objects, and famous faces). A vertex-wise significance threshold of p<0.001 was used to draw the pre-constrained ROIs.

### ROI aggregation across participants

Below we detail the procedure for establishing the probabilistic group definition for all regions of interest (scene perception areas: OPA, PPA, MPA; place memory areas: LPMA, VPMA, MPMA).

Individual participants exhibit a wide spectrum of activation extent and magnitude during the localizer tasks. To ensure that that these differences did not skew our results and that all individuals contributed equally to the probabilistic parcellation, we constrained the large ROIs drawn from the GLM to the subset of the top 800 most selective surface vertices in each region in their respective task. These vertices had the maximum difference between the category of interest compared to other categories in the task of interest (e.g., in perception, the vertices with the largest difference between scene perception vs face images in Datasets 1&2). If the region of interest had fewer than 800 vertices, all vertices were taken.

We chose a fixed number of vertices for all participants and regions to ensure that no participant biases the overall spatial extent of the final parcels. Since all participants contribute the same number of vertices, the parcel extent and peak of overlap reflect the pattern across the whole group. Using a percentage of vertices causes spatial extent of the parcel to overrepresent participants who have larger ROIs. Because the goal of generating the parcels was to provide a representative spatial location to help guide ROI drawing, we considered it essential that the parcels reflect the overall pattern of the constituent participant’s regions’ spatial extent in this unbiased way. Across subjects included in the parcels, the mean ROI size was greater than 1000 vertices (mean±std ROI size – MPA: 1135±891, PPA: 2597±880, OPA: 2181±1161, MPMA: 3444±1.304, VPMA: 2147±624; LPMA 3039±1053), and most regions had greater than 800 vertices for most participants (Figure 1).

**Figure 1.**
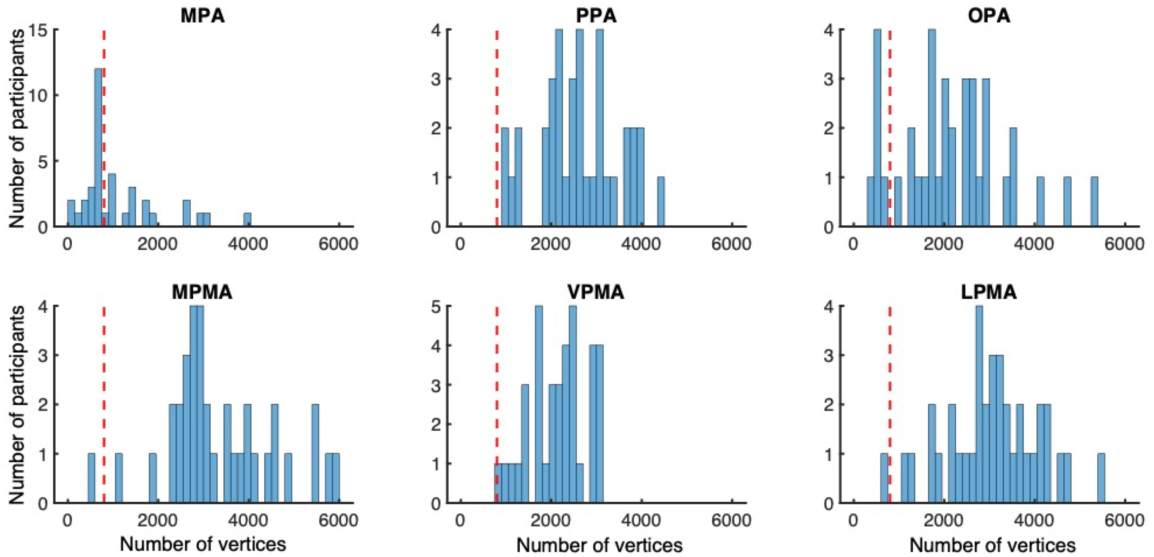
In most cases, participants had greater than 800 vertices in the hand drawn ROI prior to constraining. Histograms show the size of the hand drawn region for all participants included in the final parcellation (N=37). Red dotted line marks 800 vertices. The mean number of vertices for all regions was greater than 800, and only MPA had a significant proportion of participants with fewer than 800 vertices (median =746 vertices).

The size of the constrained region (800 vertices) was chosen to balance the size of the regions across subjects, the desire to have the most selective vertices represented, and the desire for contiguity across the vertices. Our prior publications have considered the top 300 most selective vertices (Steel et al., 2021, 2023), but this constraint has generally resulted in incontiguous sets of vertices. This led to the adoption of the new, larger set of vertices for parcel definition.

We refer to the top 800 most selective vertices as the individualized regions of interest. We aggregated these individualized regions of interest within Datasets 1, 2, and 3 for comparison. We also aggregated across all datasets to form the final probabilistic group parcel definition. We refer to these final probabilistic overlap maps of the individualized regions of interest as “parcels”.

### Comparison across parcels across datasets

We evaluated the success of our parcel definitions for the scene perception and place memory areas by comparing the location of the parcels across Datasets 1-3. For these analyses, we focused on Dataset 1 as the basis for comparison for two reasons. First, Dataset 1 was originally used to establish the place memory areas, and we wanted to confirm that this original definition was consistent. Second, Datasets 2 had the largest number of participants to serve as test cases for the generalization of the established parcels.

We performed three different quantifications: 1) overlap between group-level probabilistic maps across datasets, 2) proportion of individual constrained ROIs captured by the group parcel across tasks (i.e., how much did individual participants’ constrained ROIs from Dataset 2 & 3 overlap with the group parcel from Dataset 1), and 3) whether the individual participants’ peak of selectivity was captured by the group parcel across tasks (i.e., did the individual participants’ peak of selectivity from Dataset 2 & 3 fall within the group parcel from Dataset 1). These metrics provide a comprehensive view of the generalization of the parcels in independent data.

### Shift between scene perception and place memory selectivity

In addition to comparing the parcels across datasets, we also compared the spatial dissociation between the scene perception and place memory areas in all participants. We compared the weighted center-of-mass of the scene perception and place memory areas using the individualized regions of interest from Datasets 1 & 2 (using the methods described in (Steel et al., 2021)). Specifically, we calculated the center of mass of the individualized scene perception and place memory parcels, with the spatial position of each vertex weighted by its selectivity magnitude (i.e., activation for scene versus face perception or place versus face memory).

### Comparison with other scene perception area parcellations

Several groups have published publicly available parcels of the probabilistic locations of PPA and OPA, defined by contrasting activation during place images compared to other categories using different localizer stimuli and scan parameters (Julian et al., 2012; Weiner et al., 2018; Rosenke et al., 2021). To determine how our parcels compare with existing parcels, we systematically compared the location or our probabilistic parcels with the location of these previously established parcels. Below we briefly describe the data yielding the parcels in the previous studies (paraphrased from their methods sections) for convenience of the readers.

#### Probabilistic PPA (Weiner et al., 2018)

12 participants underwent two runs of a functional localizer task from (Stigliani et al., 2015), in which they viewed gray-scale natural images of faces (adult and child faces), scenes (corridors and houses), bodies (limbs and headless bodies), characters (numbers and pseudowords), and objects (cars and guitars). Stimuli were shown in 16 second blocks, and images from each category were on the screen for 1 second (1Hz presentation rate). Participants were instructed to fixate on a red dot during the task while performing an oddball task in which they pressed a button whenever they detected phase scrambled images, which occurred 0-2 times per block. The total scan duration was not given in this study.

PPA was defined in each participant based on the contrast of scenes > faces, objects, places, bodies, and characters with a voxel-wise threshold of t > 3. The full extent of each participant’s ROI was used to generate the overlap map.

#### VisF atlas (Rosenke et al., 2021)

19 participants underwent three runs of a multicategory localizer to define visual areas in lateral and ventral occipitotemporal cortex. This study used the same localizer task and stimuli used to define the probabilistic PPA by Weiner and colleagues, but with a different timing (Stigliani et al., 2015). Eight stimuli of one of the five categories (characters (pseudowords and numbers), bodies (whole bodies and limbs), scenes (houses and corridors), faces (child and adult), and objects (cars and instruments)) were presented in each miniblock design, each mini-block holding a duration of 4 s. Participants performed an oddball task while fixating, where they indicated the presence of a phase scrambled image using a button press. Each run consisted of 150 MRI volumes (total duration 300s).

OPA and PPA were defined by contrasting activity during scene blocks compared with all others (t > 3). The full extent of each participant’s ROI was used to generate the overlap map.

#### Julien parcels (Julian et al., 2012)

35 participants underwent four runs of fMRI scanning on a dynamic multicategory localizer task. Stimuli were three second movie clips of children’s faces, children’s bodies, objects (children’s toys), scrambled objects, and pastoral scenes used in previous studies (Pitcher et al., 2011). Within each imaging run, participants saw two blocks of each category, where six videos clips were shown (block duration = 18s). Three 18 second rest blocks were included at the beginning, middle, and end of the run, where the screen alternated between different full-screen colors once every three seconds. Each run consisted of 117 volumes (total duration 234 seconds). No task details (e.g., 1-back or oddball) were detailed in the methods section.

#### Statistical analysis

Statistics were calculated using MATLAB code (Version 2022a, MathWorks). Data distributions were assumed to be normal, but this was not statistically tested. Individual data points are shown. Raleigh’s tests were used to assess the consistency of the direction of the shift from perception to memory. Otherwise, paired t-tests were used where indicated. Alpha level of p < 0.05 was used to assess significance, and Bonferroni correction was applied where appropriate.

## Results

The current study had four aims. First, we wanted to establish the localization of the place memory areas across multiple datasets with different data acquisition parameters and experimental design considerations. Second, we wanted to show the consistent topographic extent and location of the place memory areas with respect to their relationship with the scene perception areas at an individual subject level. Third, we wanted to compare the location of the place memory and scene perception areas to other established parcels involved in scene perception at a group level (Julian et al., 2012; Weiner et al., 2018; Rosenke et al., 2021). Finally, we wanted to contextualize the place memory and scene perception areas’ locations within the cortical organization more generally, including their anatomical locations, and their location compared to large scale cortical networks and gradients. In addition, we have released a publicly available parcel for the scene perception and place memory areas for general use that can be downloaded from https://osf.io/xmhn7/.

We defined the place memory areas in individual participants by comparing fMRI activity when participants recalled personally familiar places compared with other categories. In Dataset 1 & 2, we contrasted activation when participants recalled places compared with familiar people’s faces. In Dataset 3, we compared activity when participants recalled places versus multiple other categories (places versus faces, objects, bodies, and famous faces).

In all Datasets, we defined an initial region of interest for each participant on their lateral, ventral and medial surfaces, which was hand drawn based on a contrast of t > 3.3 (p = 0.001, uncorrected) for people versus place memory (place memory localizer) or viewing scene versus face images/videos (scene perception localizer). This hand drawn region of interest was further constrained to the top 800 most selective surface vertices as each participant’s “individualized region of interest”. These individualized regions of interest served as the basis for comparing across datasets.

### The place memory areas’ location is consistent across datasets

We began by comparing the location of the most probable location of the place memory areas across our datasets. Within each dataset (datasets 1-3), we aggregated all participants’ individualized regions of interest to derive a probabilistic surface map. In this map, vertices were assigned the probability that they were located within the place memory areas across all individuals in the dataset. We refer to these probabilistic areas as parcels(Rosenke et al., 2021). The probabilistic maps for each parcel and their overlap are shown in Figure 2.

**Figure 2.**
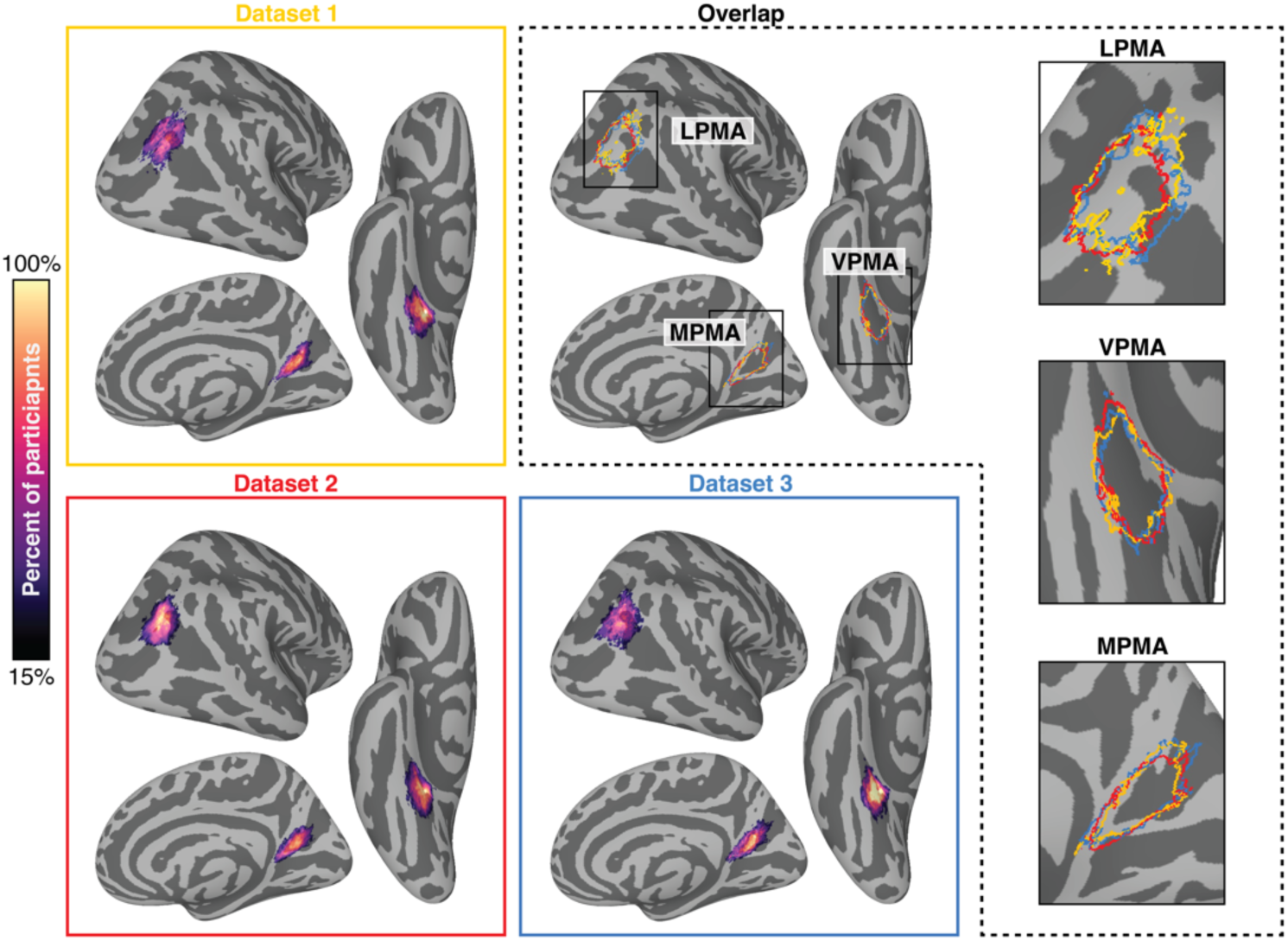
The most probable location of the place memory selective activity is consistent across datasets. The most probable location of the lateral, ventral, and medial place memory areas (LPMA, VPMA, MPMA) was based on the intersection of the top 800 most selective surface vertices for each participant. Probabilistic maps show vertices where greater than 15% of participants were represented. The extent of these regions for all datasets are outlined in the upper right panel (Dataset 1: Yellow, Dataset 2: Red, Dataset 3, Blue). The boxed portion of the image is enlarged in the insets (right).

We observed a high degree of overlap in place memory parcel’s location across the datasets, despite that the datasets considered different data acquisition, processing methods, and task conditions. Qualitatively, the spatial location of the parcels was highly similar, as was the general shape of the cluster. This confirms that the place memory area activation is highly reproducible, which suggests that these regions are a general feature of cortex and do not depend on specific design or acquisition/processing methods. We explore the location of the place memory parcels in greater detail below.

### Place memory area parcels capture individualized ROIs across datasets

Having confirmed that we could reproduce the most probable location of the place memory areas across datasets, we next quantified how well the probabilistic parcels captured the variability of the place memory region of interests’ locations in independent participants. We asked two questions: 1) would the probabilistic parcels capture the location of the most-selective vertices from individual-participants from a separate dataset, and 2) how well do the parcels capture the spatial extent of the individualized regions of interest from an independent dataset? Answering these questions will establish whether the most probable location of the parcels is an adequate description of the topography of these functionally defined areas.

For these analyses we used Dataset 1 as the basis for comparison, and we restricted the probabilistic parcels from Dataset 1 with greater than 15% of participants. We used Dataset 1 as the basis for comparison for two reasons. First, we used Dataset 1 to establish the location of the place memory areas in our original study (Steel et al., 2021), so this analysis would confirm that our original definition would generalize to new data. Second, Datasets 2 and 3 were more appropriate test datasets. Because Dataset 2 had the most participants, it provided the most opportunities to test the generalization of the parcel to new individuals. In addition, because Dataset 3 used a different experimental design, it provided an opportunity to test whether the location of the individualized regions of interest depended on the conditions used to localize these areas. Importantly, three subjects from Dataset 1 were also included in Dataset 3; however, because Datasets 1 and 3 used different conditions and came from separate testing sessions, they are still independent. Results were comparable when these participants were removed from the analysis of Dataset 3.

Overall, the place memory parcels derived from Dataset 1 generalized to individual participants from Datasets 2 and 3. Across all of the memory areas, the parcels from Dataset 1 captured the surface vertex with the greatest selectivity to place memory in the majority of participants in Dataset 2 (Left hemisphere – MPMA: 78%, VPMA: 87%, LPMA: 87%; Right hemisphere – MPMA: 78%, VPMA: 91%; LPMA: 100%) and Dataset 3 (Left hemisphere – MPMA: 80%, VPMA: 100%, LPMA: 60%; Right hemisphere – MPMA: 80%, VPMA: 100%; LPMA: 80%) (Figure 3A-B). Qualitatively, most maxima that fell outside of the parcel were very close to parcels’ edges (Figure 3A). So, it is possible that these peaks could be captured by the parcel if a more liberal threshold were used.

**Figure 3.**
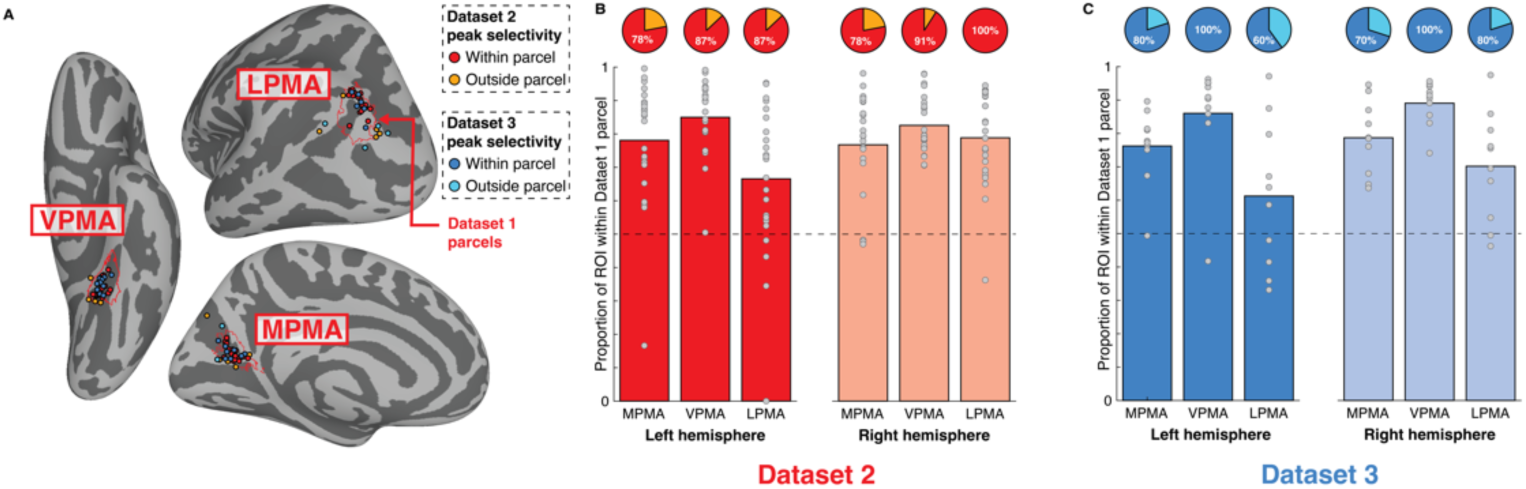
Location of place memory areas is consistent across datasets. A. Location of the peaks in place memory area parcels (red outline) compared with the individual participant peaks in place memory selective activity from an independent set of participants using di[erent fMRI processing and analysis methods (Dataset 2, red) and experimental design (Dataset 3, blue). Participants with peaks within the parcel are shown in red/blue, peaks outside the parcel are shown in yellow/cyan for Datasets 2 and 3, respectively. Only the left hemisphere is shown. These data are summarized as pie charts for both hemispheres in panel B and C. B and C. Overlap between the place memory area parcels with individual participant defined place memory area ROIs (top 800 memory selective surface vertices) from Dataset 2 (B) and Dataset 3 (C). Pie charts show the percentage of participants with peak selectivity contained within the group parcel. Dotted line indicates 50% overlap (400 vertices).

Likewise, the participants’ individualized regions of interest from Dataset 2 and 3 (top 800 most selective vertices) overlapped substantially with the parcel from Dataset 1 (Figure 3B). We observed significant overlap (> 50%) between individual participants from Dataset 2 and parcel from Dataset 1 in all regions and all hemispheres (Dataset 2: ts(23) > 3.54, ps < 0.0018). For Dataset 3, we observed significant overlap (> 50%) between individual participants in all regions in the right hemisphere (ts(9) > 3.88, ps < 0.004), and in the place memory parcel on the medial and ventral surfaces in the left hemisphere (ts(9) > 8, ps < 0.001); however, the place memory parcel on the lateral surface did not significantly capture greater than 50% of the individualized parcels across participants in Dataset 3 (t(9) = 1.62, p = 0.141). The results for Dataset 3 were similar when overlapping participants were excluded (Left hemisphere – MPMA: t(7) = 4.99, p = 0.003; VPMA: t(7) = 22.90, p < 0.001; LPMA: t(7) = 1.33, p = 0.23; Right hemisphere – MPMA: t(7) = 6.30, p < 0.001; VPMA: t(7) = 31.84, p < 0.001; LPMA: t(7) = 3.44, p = 0.014). Across both datasets, only one participant (Dataset 2) had zero vertices of their ROI confined to the LPMA parcel in the left hemisphere, which was due to the atypically anterior location of their LPMA ROI compared to other participants.

Collectively, these results show that the original place memory parcels established in the subjects from Steel et al., 2021 generalize to new datasets with different data acquisition, processing, and experimental design choices. This suggests that the place memory parcels from one dataset are adequate descriptors of these areas in new datasets, opening the possibility of algorithmic region of interest definition based on these established parcels.

### Anterior shift for place memory vs. scene perception is reproducible

The place memory areas are thought to support mnemonic processes relevant to visual scene analysis, and one key piece of supporting evidence is their proximity to the scene perception areas(Steel et al., 2021). In individual studies, the place memory areas are reported to lie consistently anterior and adjacent to the scene perception areas on their respective cortical surfaces (Steel et al., 2021; Srokova et al., 2022). Next, we tested whether the anterior topographic shift for PMAs relative to SPAs is reproducible across participants in our three datasets. For these analyses, we considered Datasets 1 and 2, which used the same experimental design. The results from Dataset 3 will be reported separately in a future publication.

We addressed the relative location of the place memory and scene perception areas in two ways. First, in Datasets 1 and 2, we considered the most probable location of the place memory areas compared to the most probable location of their functionally paired scene perception areas in the same group of participants. Given the similarity of the place memory areas’ locations in Datasets 1 and 2 described above, we combined the individualized regions of interest from both Datasets. We then compared topography of the scene perception and place memory area parcels constructed from these combined datasets, including the location of the maximum probability. To evaluate overlap, we thresholded the probabilistic place memory and scene perception parcels to represent at least 33% of participants, a threshold that is consistent with prior publications (Julian et al., 2012; Weiner et al., 2018; Rosenke et al., 2021). The peak probability for the place memory and scene perception parcels are shown in Table 2).

**Table 2.**
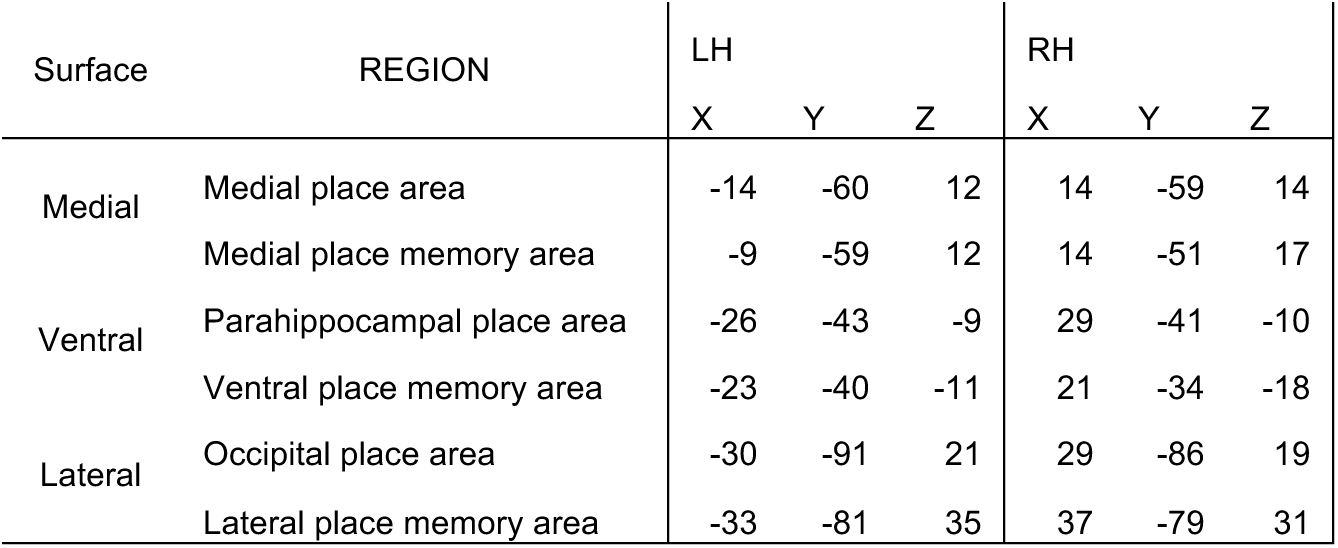
Peak probability for perception and memory parcels across cortical surfaces. Coordinates are in MNI space referenced to the MNI-N27 atlas.

We found that the place memory parcel was located anterior to the scene perception area parcel on all cortical surfaces (Figure 4). On the lateral surface, we observed an anterior and dorsal shift of the place memory parcel compared with scene perception, and we saw a relatively small amount of overlap between the memory and perception parcels (Percent of LPMA shared – Left hemisphere = 4.4%; Right hemisphere shared = 8.4%). On the ventral surface, we observed a nearly veridical anterior shift of place memory compared with scene perception and approximately half of the memory parcel was shared with the perception area parcel (Percent of VPMA shared – Left hemisphere = 41.3%; Right hemisphere = 42.9%). On the medial surface, we observed an anterior and dorsal shift in the peak probability between the parcel perception and memory parcels, but these parcels had a high degree of overlap (Percent of MPMA shared – Left hemisphere = 35.6%; Right hemisphere = 61.6%; note that the small amount of LPMA shared is due to the relatively small size of left MPA at an overlap of 33%). These results confirm the spatial dissociation between the place memory and scene perception areas on the lateral surface but suggest a less strong distinction between the memory and perception responses on the medial surface.

**Figure 4.**
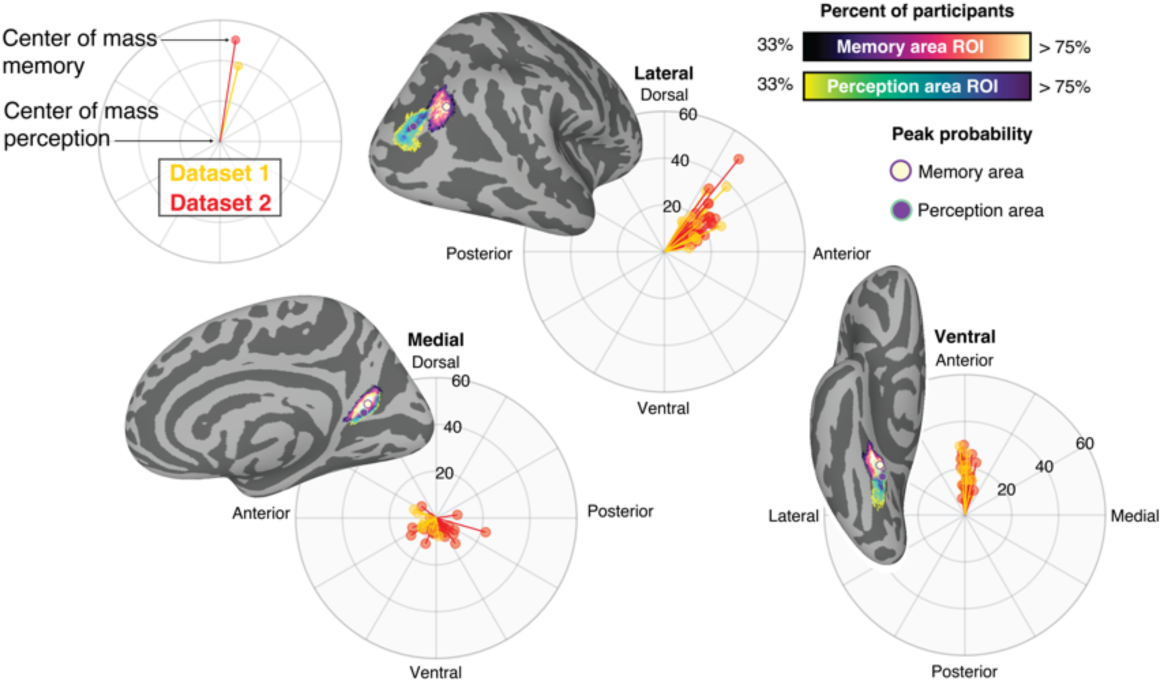
Place memory areas are consistently anterior to scene perception areas. Brain images depict the most probable location of the place memory (yellow) and scene perception (purple) areas on the brain’s lateral, medial, and ventral surfaces based on the overlap of the top 800 most selective vertices for each task across all participants. Rose plots depict the shift in the weighted center of mass oof place memory compared to scene perception activation in individual participants, split between Datasets 1 and 2 (yellow and red, respectively). This individualized analysis revealed a consistent shift of place memory compared to scene perception activity on all surfaces (Raleigh’s tests: zs > 7.6, ps < 0.001). This shift was anterior and dorsal for the lateral surface, anterior on the ventral surface, and ventral on the medial surface. Note that rose plots orientation is aligned to the respective brain surface plot. Probability maps for parcels are thresholded at 0.33 overlap across participants, consistent with prior work establishing probabilistic locations of functional areas in occipito-temporal cortex(Julian et al., 2012; Wang et al., 2015; Weiner et al., 2018; Rosenke et al., 2021).

The prior analysis shows an anterior shift of scene perception and place memory activity at the group level, but it does not capture the consistency of the shift at the individual participant level. We tested the consistency of this shift in a second analysis by comparing the location of the weighted center of mass of each participant’s individualized perception and place memory area regions of interests for each cortical surface (Steel et al., 2021). This analysis considers the degree of selectivity for the category of places/scenes versus people/faces during memory and perception, and therefore provides a sensitive measure of activation shifts between conditions in regions with a high degree of intersubject variability. Note that this analysis has been previously published using Dataset 1 with a different ROI definition method (Steel et al., 2021). Here, we replicate that analysis using a new ROI definition (top 800 most selective scene perception and place memory area vertices), and we extend that analysis by considering a new set of participants from Dataset 2. A comparison between scene perception and place memory area activity in Dataset 3 will be presented elsewhere. The results were consistent across hemispheres, so we report the average results across hemispheres.

With this approach, we observed a systematic shift of place memory compared with scene perception activation on the lateral and ventral surfaces of the brain (Figure 4). In Datasets 1 and 2, on both lateral and ventral surfaces, the individualized weighted center of mass was significantly anterior to the scene perception area (Dataset 1 – Lateral: t(13) = 10.22, p < 0.001; Ventral: t(13) = 10.11, p < 0.001; Dataset 2 – Lateral: t(22) = 15.94, p < 0.001; Ventral: t(22) = 12.17, p < 0.001). On the lateral surface, the angle of this shift was anterior and dorsal, and it highly consistent across individuals (Dataset 1 – mean±s.d. shift distance = 20.6 ±8.4 mm., Raleigh’s test: z = 13.05, p < 0.001; Dataset 2 – mean±s.d. shift distance = 21.4±9.6 mm, Raleigh’s test: z = 22.24, p < 0.001). On the ventral surface, the shift was largely anterior, with little medial-lateral variation (Dataset 1 – mean±s.d. shift distance = 17.5±7.5 mm, Raleigh’s test: z = 13.86, p < 0.001; Dataset 2 – mean±s.d. shift distance = 16.9±7.8 mm, Raleigh’s test: z = 22.71, p < 0.001). This replicates the anterior shift previously reported (Steel et al., 2021) using this new individualized ROI definition. Further, this shows that the magnitude of the shift replicates to a larger sample of participants collected using different fMRI acquisition and processing methods.

On the medial surface, the displacement of the place memory compared with scene perception activity was less pronounced. We found a significant anterior shift of the place memory compared with scene perception activation on the medial surface in Dataset 1 (t(13) = 2.47, p =0.03) along a consistent anterior and ventral angle (Mean±s.d. shift distance = 6.22±3.0 mm; Raleigh’s test: z = 8.74, p < 0.001), which replicated our prior result with the new individualized region of interest definition (Steel et al., 2021). However, we did not observe an anterior shift on the medial surface in Dataset 2. While the activity was shifted (Mean shift distance = 7.8±6.2 mm; Raleigh’s test: z = 7.76, p < 0.001), the shift of memory compared to perception was more ventral (t(22) = 5.90, p < 0.001) rather than anterior (t(22) = 0.79, p = 0.43). Note that the ventral shift of place memory compared with scene perception activity is not consistent with the most probable location of the cluster reported above, where the most probable location of the scene perception area was ventral and posterior to the most probable location of the place memory area. It is likely that the additional information provided by using magnitude of selectivity to weight the center of mass for this analysis led to this discrepancy.

Taken together, these results replicate prior work investigating the difference in topography between scene perception and place memory activation in posterior cerebral cortex(Steel et al., 2021). Specifically, the anterior shift for place memory vs. scene perception is most prominent on the lateral surface and ventral surfaces, as compared with the medial surface. This difference between cortical surfaces could belie a difference in the functional distinction between the perception and memory areas across the cortical surfaces (see Discussion).

### Location of the place memory areas compared to established scene perception area definitions

The parahippocampal place area (PPA) and occipital place areas (OPA) are critical parts of the scene perception system. These regions can be reliably localized by contrasting brain activity to images of scenes compared to images of other categories like faces, bodies, or objects; however, the precise contrast used varies between research groups. In an effort to standardize the definition of these areas, several groups have published publicly available parcels of the probabilistic locations of PPA and OPA, defined by contrasting activation during place images compared to other categories using different localizer stimuli and scan parameters (Julian et al., 2012; Weiner et al., 2018; Rosenke et al., 2021). How do the place memory and scene perception parcels defined in this study compare with the probabilistic locations of parahippocampal place area and occipital place areas from these independent functional parcels? Specifically, do the place memory areas fall anterior to the probabilistic location of PPA and OPA as defined by these independent datasets? For these analyses, we thresholded our probabilistic parcels to at 33% overlap across participants, consistent with prior work establishing probabilistic locations of functional areas in occipito-temporal cortex(Julian et al., 2012; Wang et al., 2015; Weiner et al., 2018; Rosenke et al., 2021).

We compared the location of the place memory and scene perception area parcels to three different publicly available group-defined parahippocampal place areas and two different occipital place areas. First, for parahippocampal place area, we considered a probabilistic definition of this region from Weiner and colleagues (henceforth wPPA) (Weiner et al., 2018). Second, for both parahippocampal place area and occipital place area, we considered an atlas of functionally-defined category selective areas, known as the visf atlas (Rosenke et al., 2021), that includes a scene selective area on the ventral and lateral surfaces (COS-places and TOS-places, respectively). Third, we considered the parahippocampal place area and occipital place area from Julian and colleagues (Julian et al., 2012).

In sum, we found that the peak of both our ventral and lateral place memory parcels fell anterior to the peak of every other previously published PPA and OPA/TOS parcel, respectively (Figures 5-7). With reference to the wPPA parcel from Weiner and colleagues (Weiner et al., 2018), we found that the peak of our ventral scene perception parcel corresponded closely to the peak location of the wPPA (Figure 5) (Weiner et al., 2018). While our scene perception parcel extended posteriorly beyond the wPPA, the maximum probability was well aligned between these areas. Importantly, the place memory parcel extended anteriorly beyond the wPPA, and the peak in probability of the place memory parcel was anterior to peak in probability from the wPPA. This suggests that the scene perception parcels we defined are well aligned with this definition of the parahippocampal place area (as previously shown) (Steel et al., 2021).

**Figure 5.**
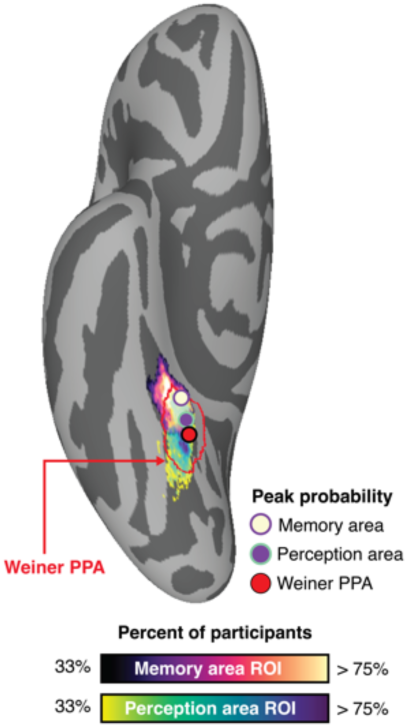
Location of most probable location of the place memory and scene perception parcels compared parahippocampal place area defined by Weiner and colleagues (red outline) (Weiner et al., 2018). Place memory area parcels combine the most probable locations of participants in Datasets 1 and 2 (33% of participants). Probable location data is reproduced from Figure 4.

Second, we considered the visf atlas (Rosenke et al., 2021), which contained a ventral and lateral scene selective area analogous to the parahippocampal place area and occipital place area (referred to as COS-places and TOS-places); notably COS-places from has a significantly greater anterior extent compared to the wPPA (Rosenke et al., 2021). We found that COS-places extended anteriorly beyond our ventral scene perception area parcel (Figure 6). Consequently, we observed more overlap between the place memory area parcel with COS-places. TOS-places also had a greater anterior extent than the lateral scene perception area parcel, and therefore also overlapped more with the place memory parcel. We found that the TOS-places region of interest overlapped with the peak of probability of both the scene perception and place memory parcels on the lateral surface. Importantly, the place memory parcels were still anteriorly shifted compared to COS-places and TOS-places, which supports our conclusion that the place memory parcels are distinct from functionally localized visual areas.

**Figure 6.**
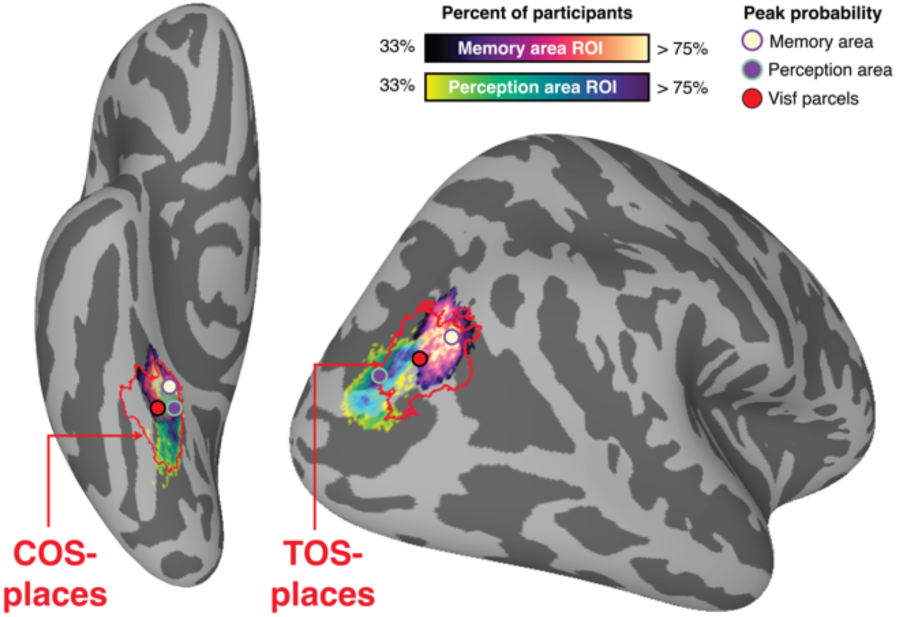
Location of most probable location of the place memory and scene perception parcels compared to COS-places and TOS-places defined in the visf atlas (red outline) (Rosenke et al., 2021). Place memory area parcels combine the most probable locations of participants in Datasets 1 and 2 (33% of participants). Probable location data is reproduced from Figure 4.

Third, we considered the parcels from Julian and colleagues (Julian et al., 2012), which contained parahippocampal place area and occipital place area atlas regions of interest (henceforth jPPA and jOPA, respectively). Interestingly, we found mixed topographic relationship between our scene perception and place memory area parcels and the Julian regions of interest (Figure 7). Although we found that the jPPA captured the maximum probability of our ventral scene perception parcel, our parcel had a larger anterior and posterior extent, and we observed relatively little overlap between the jPPA and our place memory parcel. On the lateral surface, the jOPA was shifted anteriorly compared to our lateral scene perception parcel. Like TOS-places from the visf atlas, the jOPA overlapped with the maximum probability of both our scene perception and place memory parcel. However, the lateral place memory parcel was distinct from the jOPA: the lateral place memory parcel had a greater dorsal-ventral extent compared to the jOPA, and the lateral place memory parcel was shifted anteriorly compared with the jOPA. Thus, the Julian scene perception area regions of interest were qualitatively different in their topographic extent from the parcels defined from our data.

**Figure 7.**
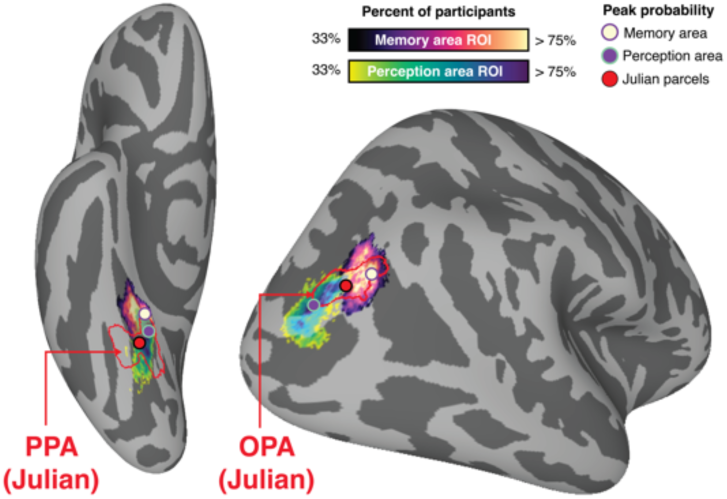
Location of most probable location of the place memory and scene perception parcels compared to parahippocampal place area (PPA) and occipital place area (OPA) regions of interest defined by Julian and colleagues (red outline) (Julian et al., 2012). Place memory area parcels combine the most probable locations of participants in Datasets 1 and 2 (33% of participants). Probable location data is reproduced from Figure 4.

### Place memory areas are at an inflection point between unimodal and transmodal cortex

The previous results demonstrated the robust localization of place memory selective areas showed the consistent topographic distinction (anterior shift) between the place memory activity compared with scene perception activity. Next, we investigated the location of the place memory and scene perception parcels compared with other proposed anatomical and functional landmarks of cerebral cortex. Comparing the locations to established landmarks will provide a more comprehensive understanding the place memory area’s position cortex more generally.

#### Glasser atlas

First, to better understand the place memory and scene perception location relative to known anatomical landmarks, we compared these parcels to the multimodal atlas from Glasser and colleagues (Glasser et al., 2016). We used the probability maps for the place memory and scene perception areas combined from Datasets 1 and 2 thresholded to consider only vertices with greater than 33% of participants represented.

The place memory and scene perception areas were largely centered on different known anatomical landmarks from the Glasser parcellation (Figure 8). On the lateral surface, the place memory parcel was situated over PGp and only minimally overlapped with other areas posterior to PGp. The place memory parcel did extend anteriorly and dorsally into PGs and intraparietal area 0 and 1. Overall the place memory parcel location may be qualitatively similar location to the tertiary sulcus slocs-v, based on visual comparison with published maps (Willbrand et al., 2024). On the other hand, the lateral scene perception parcel was situated more posteriorly, falling largely within area V3CD and posteriorly into the fourth visual area. The portion of the scene perception parcel shared with the place memory parcel did extend anteriorly into PGp and interparietal area 0, ventrally into LO1, dorsally into V3B.

**Figure 8.**
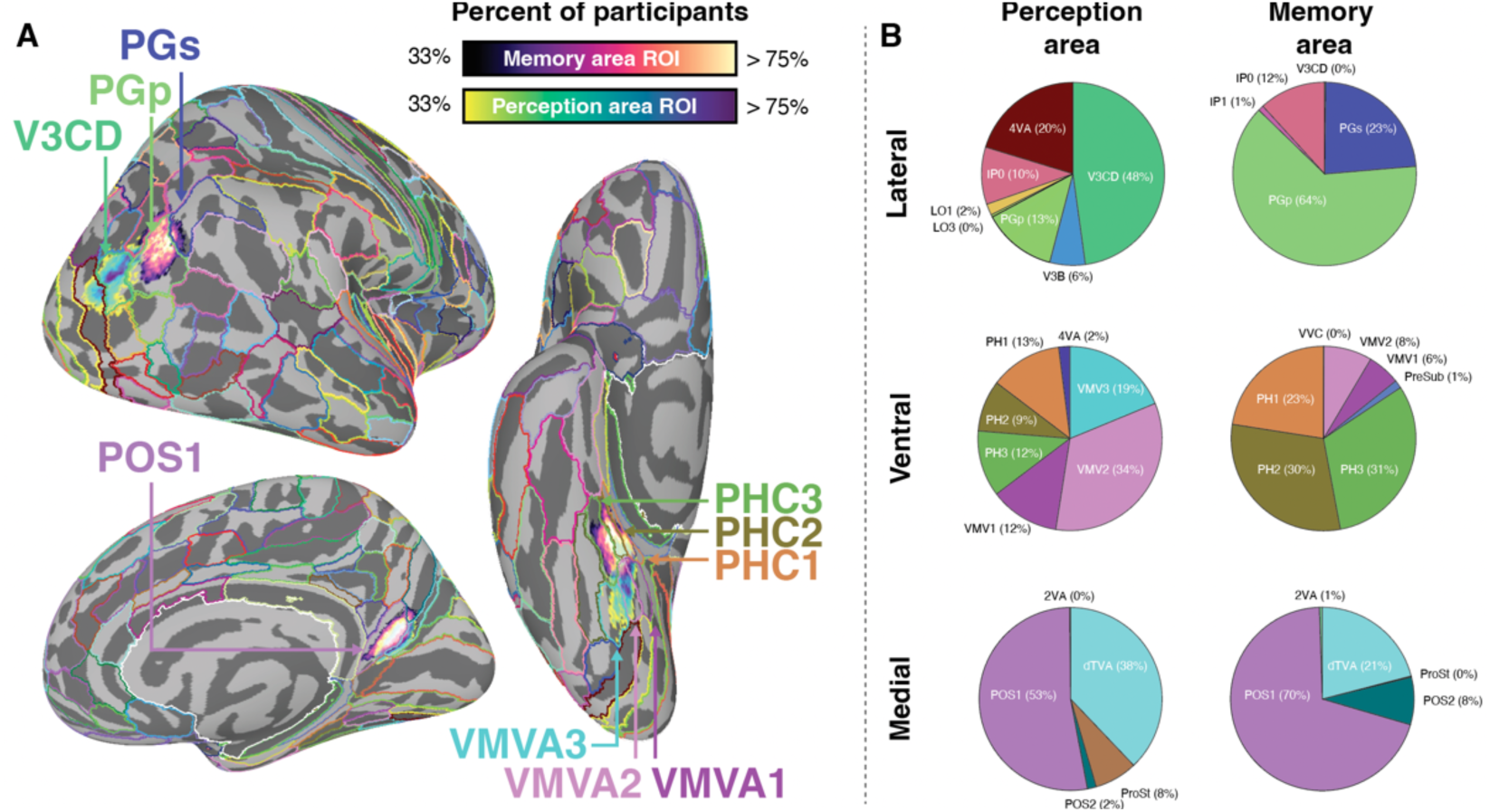
Location of most probable location of the place memory and scene perception parcels compared to the anatomical regions from the Glasser MPM atlas (Glasser et al., 2016) (Glasser 2016). A. Probabilistic parcels for perception and memory areas with Glasser atlas regions outlined. B. Pie charts showing distribution of vertices within the Glasser atlas regions. On the lateral and ventral surfaces, the peak probability of the place memory area was centered in a region anterior to the scene perception area. On the lateral surface, the maximal probability was at the boundary between PGp and PGs, while the scene perception area peak was in V3CD. On the ventral surface, the place memory area parcel was centered on PHC1-3, while the scene perception area was located primarily within the VMVA1-3. On the medial surface, both the scene perception and place memory area were largely confined within POS1. Place memory area parcels combine the most probable locations of participants in Datasets 1 and 2 (33% of participants). Probable location data is reproduced from Figure 4. Pie charts contain data from left and right hemispheres.

On the ventral surface, the place memory parcel began at the anterior portion of the ventromedial visual areas 1-3 and covered the full extent of parahippocampal areas 1-3. The medial portion of the place memory parcel crossed minimally into presubiculum. The anterior portion did not overlap with entorhinal or perirhinal cortex, and the posterior extent of the place memory parcel only minimally overlapped with the ventromedial visual areas 1-3. In contrast, the ventral scene perception parcel was situated within the ventromedial visual areas 1-3, and centered on ventromedial visual area 2. The scene perception parcel did extend anteriorly into the middle of parahippocampal areas 1-3.

On the medial surface, both the medial place memory and scene perception parcels covered the full extent of parieto-occipital area sulcus area 1 and terminated prior to retrosplenal complex. Both parcels overlapped with the transitional visual area and parieto-occipital sulcus area 2. Neither the place memory or scene perception parcel extended anterior-dorsally into area 23 A/B or area 7M.

#### Retinotopic maps (Wang atlas)

Retinotopic maps, topographic recapitulations of the retina based on voxels’ preferential responses to stimulation of portions of the visual field, are a hallmark of visual cortical organization. Although the visual field preferences are shared between retinotopic maps, many of the retinotopic maps have dissociable roles in visual processing (e.g., the independent processing of orientation and shape in LO1 versus LO2 (Silson et al., 2013)). Understanding the location of the scene perception and place memory parcels relative to these retinotopic maps will help understand these areas’ locations compared to these well-characterized sub-regions of classically visually responsive cortex. To this end, we compared the most probable location of these parcels with the most probable location of the retinotopic maps using the atlas from (Wang et al., 2015).

We found that the scene perception parcel largely fell within the retinotopic maps, while the place memory parcel was generally anterior to these maps, consistent with their respective roles in perception and memory (Figure 9). On the ventral surface, the peak of scene perception area probability fell within map PHC2, and the parcel extended posteriorly as far as map VO1. In contrast, the peak probability of the ventral place memory area parcel was anterior to map PHC2. The place memory area parcel did extend posteriorly into the PHC2 parcel, but much of the parcel was anterior to the map.

**Figure 9.**
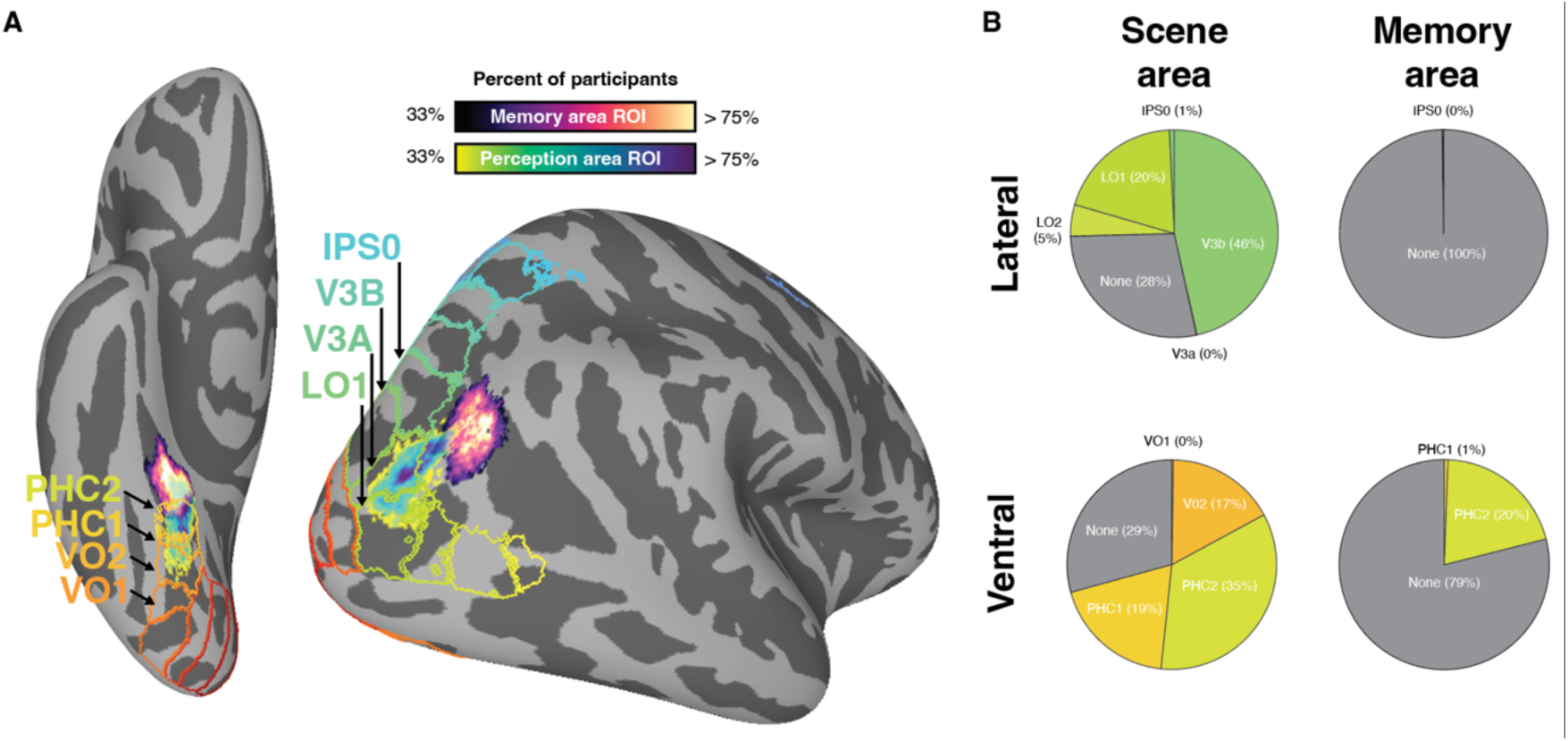
The place memory areas fall outside of the retinotopic maps, while the scene perception areas fall largely within these maps. A. Probabilistic parcels for lateral and ventral scene perception and place memory areas with retinotopic maps overlaid. Retinotopic maps were defined using a maximum probability projection from a standard atlas (Wang et al., 2015). B. Distribution of vertices within each retinotopic map combined across hemispheres. Place memory area parcels combine the most probable locations of participants in Datasets 1 and 2 (33% of participants). Probable location data is reproduced from Figure 4.

On the lateral surface, the location of the perception and memory parcels followed a similar pattern to the ventral surface. Most of the scene perception parcel fell within the retinotopic maps LO1, V3A, and V3B, and the peak of scene perception parcel probability fell within V3A. On the other hand, the place memory parcel was anterior to these maps, nestled within the space not considered classically visually responsive between maps LO2, LO1, V3A, V3B, IPS0, and IPS1.

Taken together, these results show that the place memory parcels are largely anterior to classic retinotopic maps. This is generally consistent with the assumed dissociation with their functional roles. Notably, recent work has shown that the place memory areas have nontraditional visual responses, including systematic negative BOLD responses to stimulation of positions on the retina (negative retinotopic coding) (Angeli et al., 2024; Steel et al., 2024b, 2024a), consistent with much of the default mode network of the brain (Szinte and Knapen, 2020; Christiaan Klink et al., 2021). The difference in visual coding between the scene perception and place memory areas adds to the evidence for their distinct roles in cognition.

#### Large-scale cortical networks (Yeo HCP 15)

Within human cognitive neuroscience, there is a growing consensus that the coordinated activity across large-scale distributed networks of brain areas underpins cognitive processes like external attention, episodic projection, social processing, and language comprehension and production (Fox et al., 2005; Buckner et al., 2011; Power et al., 2011; Braga and Buckner, 2017; DiNicola et al., 2020; DiNicola and Buckner, 2021; Du et al., 2024; Steel et al., 2024a). Establishing where the place memory and scene perception parcels are located compared to these distributed networks will help contextualize how these regions contribute to the brain’s large-scale activity. Importantly, these distributed cortical networks can be defined within individual participants with sufficient resting-state fMRI data(Laumann et al., 2015; Braga and Buckner, 2017; Gordon et al., 2017; Gratton et al., 2018; DiNicola and Buckner, 2021; Du et al., 2024). However, it is also common to use established templates of these networks for group analysis (Buckner et al., 2011) or as priors for individualized network definition (Kong et al., 2021; Du et al., 2024). So, here we compared the most probable location of the place memory and scene perception parcels to a commonly used template from Yeo and colleagues (Buckner et al., 2011).

Overall, the scene perception and place memory parcels were associated with networks involved in external and internally oriented attention, respectively (Raichle et al., 2001; Fox et al., 2005, 2006; Steel et al., 2024a) (Figure 10). Specifically, the peak probability of the scene perception area parcels on the lateral and ventral surfaces fell within the dorsal attention network A, a network typically associated with externally oriented attention due to its activation during attentionally-demanding visual tasks (e.g., visual search) (Fox et al., 2005, 2006). On the other hand, the place memory parcels tended to fall at the boundary between the dorsal attention network B and the default network A, generally associated with high-level visual processing and episodic projection and scene construction tasks, respectively (Braga and Buckner, 2017; Dixon et al., 2017; DiNicola et al., 2020). On the medial surface, both place memory and scene perception parcels were at the posterior-ventral edge of the large default network A cluster in posterior parietal cortex.

**Figure 10.**
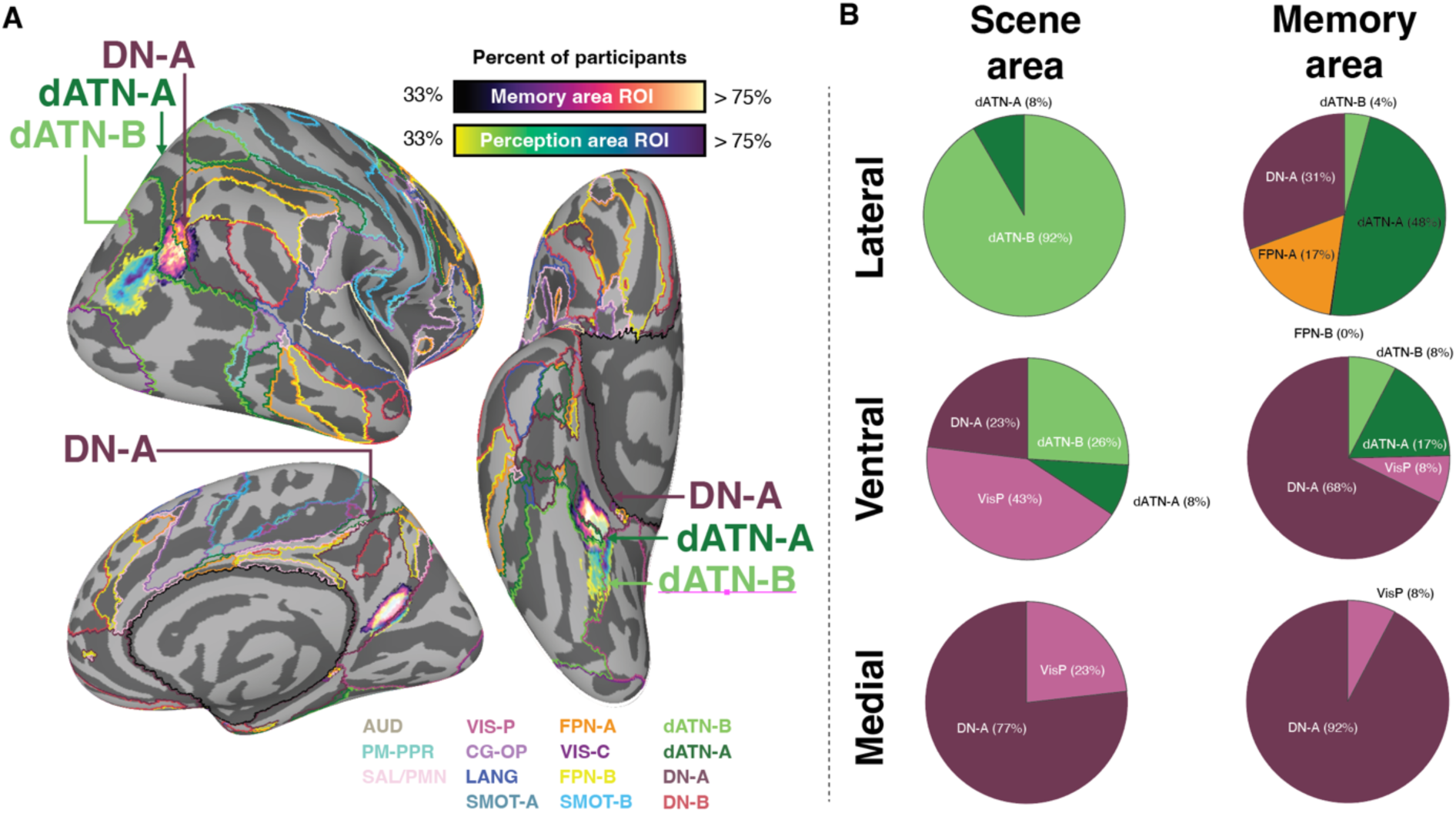
Location of most probable location of the place memory and scene perception parcels compared to the large-scale distributed cortical networks defined by Yeo and colleagues (colored outlines) (Thomas Yeo et al., 2011). A. Probabilistic parcels of the scene perception and place memory areas with the resting-state networks from Yeo and colleagues overlaid. B. Pie charts showing the distribution of vertices from each parcel within the Yeo networks. On the lateral and ventral surfaces, the scene perception areas fell largely within sensory grounded cortical networks, specifically dATN-B. On the other hand, the place memory area parcels fell within networks associated with more abstract, mnemonic processing, including the DN-A and dATN-A. On the medial surface, both the scene perception and place memory area parcels fell within DN-A. Place memory area parcels combine the most probable locations of participants in Datasets 1 and 2 (33% of participants). Probable location data is reproduced from Figure 4.

These data are consistent with the role of the scene perception area in high-level visual analysis. Moreover, they suggest that the place memory parcels may be a transitional zone between external and internally oriented neural systems that are not well captured by the canonical networks. However, this should be interpreted with caution, as the topography of these networks is more accurate when defined in individual participants (Gordon et al., 2017).

#### Principal gradient (Margulies et al., 2016)

Beyond discrete networks, cortical organization can be characterized by continuous gradients that transition from unimodal sensory/motor areas to transmodal association areas. These gradients are thought to reflect increasingly abstract and integrated information processing (Margulies et al., 2016; Huntenburg et al., 2018; Murphy et al., 2018, 2019). Given that place memory areas appear to bridge perceptual and mnemonic processes, understanding their position along this gradient could provide insight into their functional role. Using the principal gradient framework established by Margulies et al., we examined where scene perception and place memory areas fall along this unimodal-to-transmodal axis (Margulies et al., 2016). We hypothesized that scene perception areas, which process immediate visual input, would occupy a relatively unimodal position. In contrast, we predicted place memory areas would fall further along the gradient toward transmodal cortex, reflecting their role in integrating visual and mnemonic information.

For this analysis, we compared the location of the place memory and scene perception parcels with the first principal gradient published by Margulies and colleagues (Margulies et al., 2016). We restricted these parcels to surface vertices with greater than 33% of participants.

We found that the place memory and scene perception parcels were in distinct portions of the unimodal-to-transmodal axis (Figure 11), which we quantified by comparing the distributions of vertex-wise gradient score for between the perception and memory areas on each surface using two-sample Kolmogorov-Smirnov tests (lateral surface: D(3130)=0.90, p<0.001; ventral: D(3777)=0.53, p<0.001; medial: D(2819)=0.31), p<0.001. Consistent with our hypothesis, the scene perception parcels on the lateral and ventral surfaces fell largely within unimodal cortex, while the MPA parcel tended to have an elevated position on the principal gradient compared with the other scene perception areas. Intriguingly, all place memory parcels and the MPA parcel fell at the inflection point between unimodal and transmodal cortex, and the parcels spanned the dominantly unimodal to more dominantly transmodal cortical territory (Figure 11B-D). This is further support for the distinct roles of the lateral and ventral scene perception compared with MPA and the place memory areas, and specifically that MPA and the place memory areas may constitute a unique transition zone between sensory and abstract representations.

**Figure 11.**
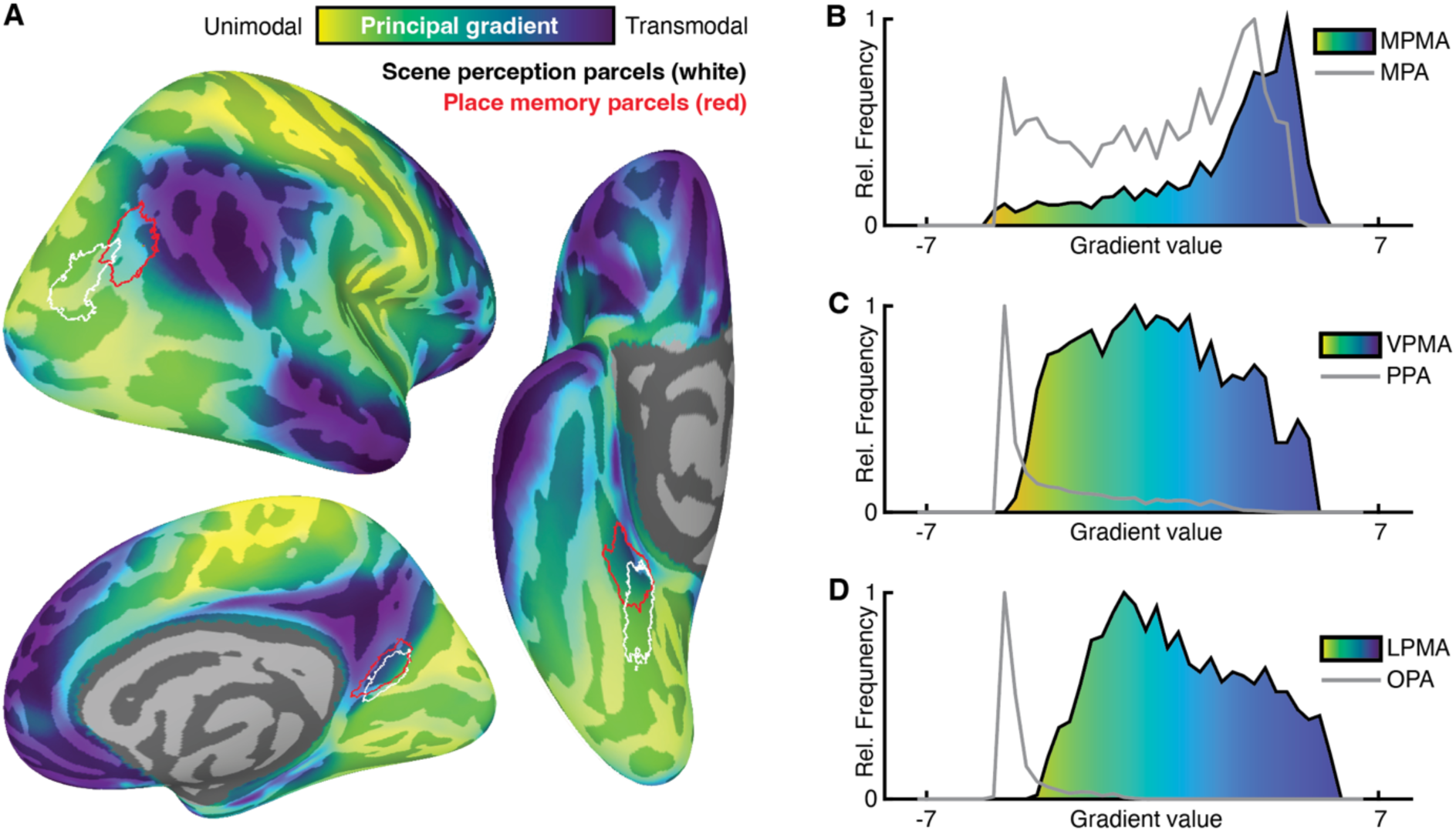
The scene perception and place memory areas are located at di[erent points along the unimodal-to-transmodal principal gradient. A. Distribution of the principal gradient on the cortical surface with scene perception and place memory area parcels overlaid. B-C. Histograms depicting the relative frequency of vertex-wise principal gradient scores for scene perception (grey) and place memory area parcels (black, colored fill). On the lateral and ventral surfaces, the scene perception parcels are largely confined to unimodal cortical territory. In contrast, the place memory parcels and the MPA parcel fall at the inflection point between unimodal and transmodal cortical territory. Principal gradient data are taken from Margulies et al., 2016(Margulies et al., 2016). Place memory and scene perception parcels were binarized probability maps containing at least 33% of participant’s individualized perception or memory area regions of interest. Histograms show data from both hemispheres.

### Place memory and scene perception parcel availability and access

Taken together, these data support the distinction between scene perception and place memory related activity on the lateral and ventral surfaces. Figure 12 shows a binarized mask of the probabilistic definition of the scene perception and place memory parcels for both hemispheres reported here (thresholded at 15% participants), which are available for download here: https://osf.io/xmhn7/. The 15% threshold was chosen because of the relative success in capturing the individual local maxima across independent datasets (Figure 3, above). Parcels are available as volume (nifti) and surface files (SUMA’s NIML standard mesh 141 density (Argall et al., 2006; Saad and Reynolds, 2012) and Gifti formatted for the fsaverage template (Fischl, 2012)). We have released a probabilistic definition of these parcels for users who wish to set their own parcel probability criteria. Finally, we have released stimulus presentation code and instructions to run the place memory localizer used in this study.

**Figure 12.**
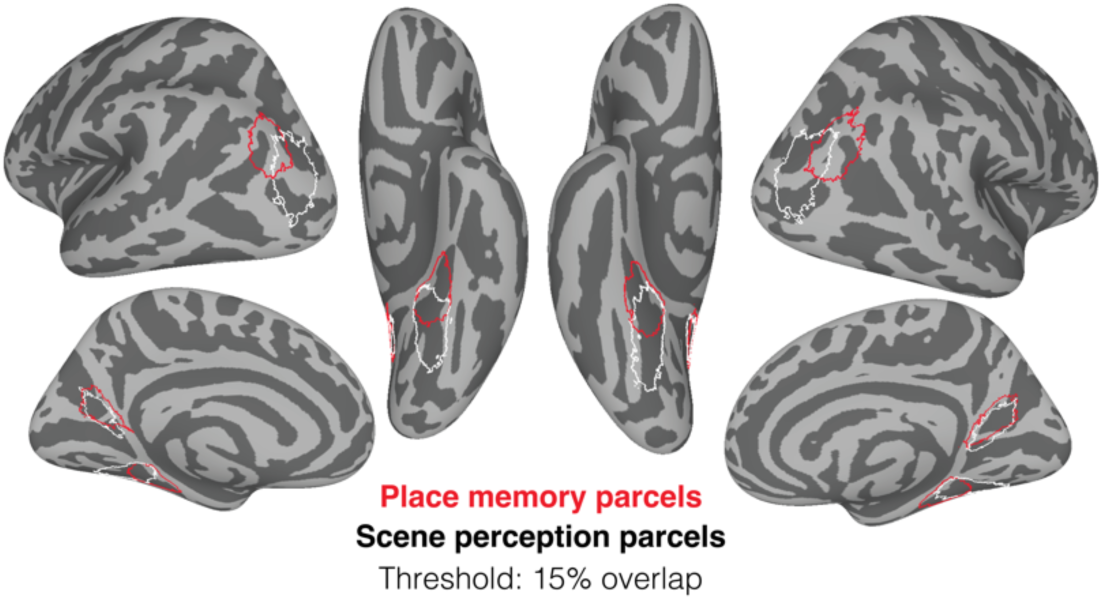
Place memory and scene perception area binarized parcels. Vertices within the parcel were present in greater than 15% of participant’s individualized perception or memory area regions of interest.

## Discussion

Here, we characterized the anatomical locations of the place memory areas, a set of brain areas that respond more when participants recall personally familiar places compared with other types of stimuli (Steel et al., 2021), across a relatively large group of participants spanning three different studies. We made three central observations. First, the place memory areas’ locations are consistent across participants and different fMRI acquisition parameters, localizer tasks, and statistical contrasts. Second, within a participant, the place memory areas lie systematically anterior to the scene perception areas on the lateral and ventral surfaces. Third, at the group-level, compared with the scene perception areas, the place memory areas are in regions associated with memory processes. Specifically, the place memory areas are immediately anterior to the retinotopic maps and previous published definitions of the scene perception areas(Julian et al., 2012; Wang et al., 2015; Weiner et al., 2018; Rosenke et al., 2021), at the boundary between Dorsal Attention Network B and the Default Network A (Buckner et al., 2011), and span the transition between unimodal and transmodal cortex(Margulies et al., 2016). These group analyses support the hypothesis that the place memory areas serve as a bridge between perceptual and mnemonic processes in cortex(Steel et al., 2021, 2023, 2024b, 2024a; Angeli et al., 2024).

### Place memory areas are consistently anterior to scene perception areas on the lateral and ventral surfaces

A striking feature of the place memory and scene perception areas is their topographic arrangement in cortex. Our initial description revealed that the place memory areas were located immediately anterior and adjacent to the scene perception areas within individual subjects (Steel et al., 2021), which was subsequently replicated by our group and others (Srokova et al., 2022; Steel et al., 2023, 2024b). The precise arrangement between perception and mnemonic areas varies by cortical surface: the trajectory between from scene perception to place memory area is dorsal and anterior on the lateral surface, anterior and ventral on the medial surface, and anterior on the ventral surface. In addition, earlier studies found a larger anterior shift for the lateral and ventral place memory areas vs. their corresponding scene perception areas, as compared with the medial areas (Steel et al., 2021; Srokova et al., 2022). Our findings here replicate these prior findings, with remarkable consistency between the Datasets. The reproducibility of this shift across participants and paradigms suggests that a neuroanatomical linking mechanism (e.g., structural connectivity) may underpin their co-localization.

Unlike the lateral and ventral surfaces, the medial place memory and scene perception areas were highly overlapping, and the shift of memory versus perception was smaller and less consistent. The findings from Dataset 1 replicate the original observation of an anterior (and ventral) shift for the center of mass of memory compared with perception on the medial surface (Steel et al., 2021). However, we found no anterior shift in Dataset 2. Instead, we observed a ventral shift of memory compared with perception in Dataset 2, but this shift was small compared with the shift on the other cortical surfaces (approximately 7mm shift on the medial versus 18 or 21 mm shift on the ventral and lateral surfaces). In both Datasets, we observed a considerably more overlap between the perception and memory areas on the medial surface compared with the lateral and ventral surfaces. As noted in the initial studies of these areas, these differences suggest that the relationship between memory and perception on the medial surface may be categorically different than the lateral and ventral surfaces(Steel et al., 2021, 2023).

What does the co-localization memory and perceptual activity for scenes on the medial surface imply about the medial scene regions’ functions? The retrosplenial cortex (RSC), an anatomical region in the posterior medial parietal cortex near the parietooccipital sulcus near MPA, receives input from, among other areas, parahippocampal cortex, suggesting that this area may be higher in the visual hierarchy compared to this area (Kobayashi and Amaral, 2003; Alexander et al., 2023). Indeed, prior work suggests that the medial scene perception area may not be involved in visual analysis, per se, like image processing related to structure identification (i.e., “this is a tower”) or path detection, associated with the ventral and lateral surfaces, respectively (Marchette et al., 2015; Julian et al., 2016; Kamps et al., 2016; Persichetti and Dilks, 2016, 2019; Bonner and Epstein, 2017; Epstein and Baker, 2019; Henriksson et al., 2019; Lescroart and Gallant, 2019; Dilks et al., 2022). Instead, the medial scene perception area may be involved with more abstract processes related to joint visuo-mnemonic representation(Epstein et al., 2007; Park et al., 2007; Baumann and Mattingley, 2010; Vass and Epstein, 2013; Marchette et al., 2014; Robertson et al., 2016; Silson et al., 2018; Berens et al., 2021). Likewise, RSC/MPA has been implicated in scene memory and navigation (Maguire et al., 1998; Epstein et al., 2007; Auger et al., 2015; Chrastil et al., 2018; Steel et al., 2019, 2021; Foster et al., 2023). RSC/MPA is more active when viewing images of familiar scenes compared with unknown scenes(Epstein et al., 2007; Steel et al., 2021), and when viewing familiar scenes the posterior medial parietal cortex represents allocentric position of that scene in the environment (Baumann and Mattingley, 2010; Vass and Epstein, 2013; Marchette et al., 2014; Nau et al., 2020). In addition, if the spatial relationship between scenes is known (e.g., they appear on opposite sides of a street), the MPA represents the spatial relationship between these viewpoints within a larger panoramic environment (Robertson et al., 2016; Berens et al., 2021). This portion of cortex also activates during mental imagery of scenes, when recalling past events, and imagining future events (Gilmore et al., 2015, 2016; Vass and Epstein, 2017; Silson et al., 2018, 2019; Barry et al., 2019; DiNicola et al., 2020). Collectively, these functions rely on mnemonic information associated with familiarity.

Taken together, these results show a qualitative distinction between the importance of memory in MPA, and they suggest that MPA could be viewed as a region further down the visual hierarchy and not a core sensory region. Consistent with this view, in our data, we observe little distinction between perceptual and mnemonic activity on the medial surface, with visual scene perception tasks activating just a small portion of this larger, memory responsive region. Therefore, we suggest that future work distinguishing between mnemonic and perceptual processing in scenes might jointly consider this scene perception area (MPA/RSC) and its mnemonic counterpart MPMA as single memory area, akin to LPMA and VPMA, and separate from scene perception areas on the ventral and lateral surface (PPA and OPA). Notably, at a finer scale, the anterior and posterior banks of the parietooccipital sulcus may be differentially involved in memory and perception (Silson et al., 2016, 2018), and it is possible that these fine-grained distinctions have been blurred at 3T resolution used in this study. Future work using high- or layer-resolution fMRI may more clearly characterize the visual selectivity and processes in the medial scene perception and place memory areas.

### Place memory areas span the transition from unimodal to transmodal cortex

Our analyses revealed the place memory areas’ intriguing position at the intersection of perceptual and mnemonic cortical systems. Rather than simply activating either sensory regions or memory-related networks like the default network(DiNicola et al., 2020; Du et al., 2024), these areas emerge at their intersection, suggesting they may serve as a functional bridge between these systems. This pattern was particularly clear in our large-scale network analyses, where both VPMA and LPMA peaked at the boundary between the dorsal attention network B and the default network A, which are associated with external and internally oriented attention respectively(Fox et al., 2005; Dixon et al., 2017; Murphy et al., 2018, 2019). Likewise, the place memory areas fell at the inflection point between unimodal and transmodal cortical territory (Margulies et al., 2016). This position allows these areas serve as a topographic link between perceptual and mnemonic systems to facilitate their interaction during complex behaviors like navigation that require continuous interplay between perception and memory.

The transitional nature of the place memory areas was further supported by their relationship to know retinotopic maps (Wandell et al., 2007; Wang et al., 2015). Our group analysis underscored that the memory areas fell at the anterior edge of the known retinotopic maps, implying that they are interfacing with retinotopic information. However, these areas may not have a map-like topographic layout typically associated with visually responsive areas. Instead, our previous work has shown that these regions have a significant proportion of negative retinotopic voxels (Steel et al., 2024b). Recent work has shown that this bivalent (negative vs. positive) retinotopic code underpins voxel-scale interactions across place memory and scene perception areas (Steel et al., 2024b), and broader perceptual-mnemonic interactions in the brain, including interactions between the default network and dorsal attention network (Steel et al., 2024a) and between the hippocampus and cortex (Angeli et al., 2024). The place memory areas’ position at the edge retinotopic cortex and their mixed response properties suggests that they may organize the retinotopic information transfer across cortical systems. However, because these results consider group-aligned data, future work with high-resolution fMRI with dense sampling individual participants will be critical to investigate these areas’ possible role in bridging perceptual and mnemonic representation and whether these transitional areas are a general motif of cortical organization.

An important contribution of this work is characterizing place memory areas within previously established anatomical atlases and functional frameworks. For example, one prominent framework of perceptual-mnemonic systems, the PMAT framework (Ranganath and Ritchey, 2012; Ritchey et al., 2015; Reagh and Ranganath, 2018, 2023; Barnett et al., 2021), posits a “medial temporal network” (MTN), a set of brain areas including the parahippocampal cortex and precuneus, bridges visual areas to the DMN and the hippocampus ((Barnett et al., 2021), see also (Andrews-Hanna et al., 2014; Gilmore et al., 2016)). Additionally, within the DMN, two subsystems – the posterior medial networks (PMN) and the anterior temporal network (ANT) – are thought to represent place (i.e., location, situation) and item information (i.e., objects and people), respectively (Ranganath and Ritchey, 2012; Ritchey et al., 2015; Reagh and Ranganath, 2018; Cooper and Ritchey, 2019; Ritchey and Cooper, 2020). Consistent with the PMAT framework, our group analysis shows that the place memory areas overlap with the MTN described by Barnett and colleagues based on their associated Glasser parcels (Glasser et al., 2016). Specifically, both the place memory areas and the MTN are associated with the parahippocampal areas 1-3 and the PGp on the ventral and lateral surface, respectively. Showing this correspondence deepens our understanding of the place memory areas broader functional profile, implicating them more directly in autobiographical memory, scene construction, and navigation (Ranganath and Ritchey, 2012; Cooper and Ritchey, 2019; Ritchey and Cooper, 2020; Barnett et al., 2021). It also suggests that the place memory areas’ visual properties and functions during memory-guided visual tasks, including their retinotopic coding (Steel et al., 2024b) their role in representing visuospatial context (Steel et al., 2023), could be integrated within the PMAT framework. More broadly, this approach of situating brain areas within multiple reference frameworks helps synthesize findings across different theoretical perspectives and methodological approaches.

### Place memory areas can be localized across diJerent datasets and protocols

Our results show that the place memory areas are localizable in a larger group of participants and when considering a broader set of categories in memory (as is common in studies of perception, e.g., (Stigliani et al., 2015; Weiner et al., 2018; Gomez et al., 2019; Allen et al., 2021; Rosenke et al., 2021)). Importantly, we found these areas in all 44 participants we scanned, and the location of the place memory areas was consistent between groups using simple two-category contrast (familiar places versus familiar faces memory recall) or multiple categories (familiar places versus familiar faces, objects, bodies, famous faces). Taken together, these results show that the place memory areas a robust feature of human functional brain organization.

Interestingly, although the place memory parcels were significantly more anterior than all previous scene perception area parcels, including the wPPA (Weiner et al., 2018), CoS-places and TOS-places region (visf atlas)(Rosenke et al., 2021), and jPPA and jOPA (Julian parcels) (Julian et al., 2012), our lateral scene perception parcel was generally more posterior than previous scene perception parcels (TOS-places and jOPA). Several methodological differences may explain this partial discrepancy. First, these prior parcels were defined by contrasting activation when participants viewed scenes versus multiple other visual categories, while we used a more targeted scenes versus faces contrast to match our place memory localizer (places versus people recall). Second, task demands differed across studies - we used passive viewing whereas the data used to define the visf atlas employed an oddball task (Rosenke et al., 2021); Julian et al. did not report task details(Julian et al., 2012). These differences in stimuli and attention demands could contribute to the observed variability in parcel location. Taken together, our results reinforce the distinction between scene perception and place memory areas on both lateral and ventral surfaces, and our publicly available probabilistic parcels complement existing resources by providing bilateral, probabilistic ROIs that capture this distinction.

## Conclusion

To summarize, understanding the neural systems that subserve perceptual-mnemonic interactions is a critical question for neuroscience. Here, we describe the anatomical location of the place memory areas (PMAs) on the brain’s lateral, ventral and medial surface, which are well poised to facilitate perceptual-mnemonic interactions for scenes. Further study of these brain areas could yield critical insights into neural processes underpinning spatial cognition and other functions that require the dynamic interplay between perception and memory. To support this, we have released the probabilistic maps and parcels defined in the large group of participants for public use (https://osf.io/xmhn7/).

## Data availability

Probabilistic parcels and fMRI localizer tasks are available from https://osf.io/xmhn7/. Other supporting data will be made available upon publication.

## Code availability

This data does not use any original code. Any additional information required to reanalyze the data reported in this paper is available from the lead contact upon request.

## Acknowledgements

This work was supported by an award from the National Institutes of Mental Health (R01MH130529) to CER. AS was supported by the Neukom Institute for Computational Sciences.

## Author contributions

AS and CER designed the research. AS, DP, BDG collected and processed the data. AS analyzed the data. AS and CER wrote the manuscript. All authors reviewed and edited the manuscript.

During the preparation of this work, the author(s) used Claude (Anthropic) to assist in revising the manuscript. After using this tool/service, the author(s) reviewed and edited the content as needed and take(s) full responsibility for the content of the publication.

